# *In silico* conformational features of botulinum toxins A1 and E1 according to the intraluminal acidification

**DOI:** 10.1101/2022.09.01.506163

**Authors:** Grazia Cottone, Letizia Chiodo, Luca Maragliano, Michel-Robert Popoff, Christine Rasetti-Escargueil, Emmanuel Lemichez, Thérèse E. Malliavin

## Abstract

Although the botulinum neurotoxins (BoNTs) are among the most toxic compounds found in nature, their molecular mechanism of action is far from being elucidated. A key event is the conformational transition due to the acidification of the interior of synaptic vesicles, and leading to the translocation of the BoNT catalytic domain into the neuronal cytosol. To investigate these conformational variations, homology modelling and atomistic simulations are combined to explore the internal dynamics of the subtypes BoNT/A1, the most-used in medical applications, and BoNT/E1, the most kinetically efficient. This first simulation study of di-chain BoNTs in closed and open states includes the effects of neutral and acidic pH. The conformational mobility is driven by domains displacements; the ganglioside binding site in the receptor binding domain, the translocation domain (HC_NT_) switch and the belt *α* helix visit multiple conformations depending on the primary sequence and on the pH. Fluctuations of the belt *α* helix are observed for closed conformations of the toxins and at acidic pH, and patches of more accessible residues appear in the same conditions in the core translocation domain HC_NT_. These findings suggest that during translocation, the larger mobility of belt could be transmitted to HC_NT_, leading to a favorable interaction of HC_NT_ residues with the non-polar membrane environment.

**Key Contribution:** The molecular dynamics simulations presented here provide a structural and functional annotation of full-length BoNTs composed by two distinct protein chains. Two different conformations (open and closed) as well as two different protonation states, corresponding to acidic and neutral pH, have been considered. Results from the present work supports a model of mobility in which the individual domains fluctuate around stable conformations and the overall structure mobility arise from relative displacements of the domains.

## 1. Introduction

The botulinum neurotoxins (BoNTs), produced by *Clostridium botulinum*, are among the most powerful toxic compounds found in nature, provoking the typical deadly flaccid paralysis of the host [1]. BoNTs are traditionally classified into seven serotypes, termed A-G [2] and more recently discovered H or FA, J and X [3,4]. Among them, BoNT/A1 subtype is the most-used toxin in medical applications. Despite the fact that all botulinum neurotoxin serotypes A-G inhibit acetylcholine release, they hijack different neuronal receptors, cleave different intracellular components of the Soluble N-ethylmaleimide-sensitive-factor Attachment protein Receptor (SNARE) machinery that underpins the fusion of neurotransmitter-containing vesicles, and exhibit different kinetics of neuron intoxication and intracellular stability [5,6]. In particular, BoNT/A1 and BoNT/E1 display quite different properties.

As for their structural architecture, the proteolytically activated BoNTs are formed by two protein chains connected by one disulfide bridge: the light chain (LC) and the heavy chain (HC). During the last fifteen years, numerous X-ray crystallographic structures of BoNTs, obtained from samples prepared as single protein chains, were determined [7–9]. Although these structures show conformational variations, they all display similar domain organization. The Figure 1A has been prepared to give an overview of this organization using as example the BoNT/A1 structure [7]: the LC (green color) contains the catalytic site, whereas HC is composed of two distinct domains: an N-terminal translocation domain HC_N_ (≃50 kDa) responsible for the LC delivery into the cytosol, and a C-terminal domain (HC_C_) (≃50 kDa), responsible for receptors binding. HC_N_ spans the belt (cyan) and the core translocation domains (HC_NT_, orange) whereas HC_C_ spans the N terminal and C terminal receptor binding domains (HC_CN_: magenta and HC_CC_: red). In more details, the catalytic domain displays an *α-β* fold. the translocation domain HC_NT_ is composed of a bundle of *α* helices and loops. HC_CN_ contains predominantly *β*-sheets arranged into a jelly roll motif and HC_CC_ folds into a *β*-trefoil. For sake of clarity we denote the extremity of HC_NT_ located closest to the disulfide bridge, marked with an asterisk in Figure 1A, as top of HC_NT_, the opposite extremity being the bottom of HC_NT_. The two long *α* helices of HC_NT_ are denoted helix 1 and helix 2 (Figure 1B). Two other sub-domains of HC_NT_ are the HC_NT_ switch, formed by three *α* helices, and located in the middle of HC_NT_ and on the other side of HC_NT_, the HC_NT_ C terminal *α* helix (Figure 1C). It should be noticed that the C terminal *α* helix, present in the BoNT/A1 structure [7], is unfolded in the X-ray crystallographic structure of BoNT/E1 [9].

**Figure 1.**
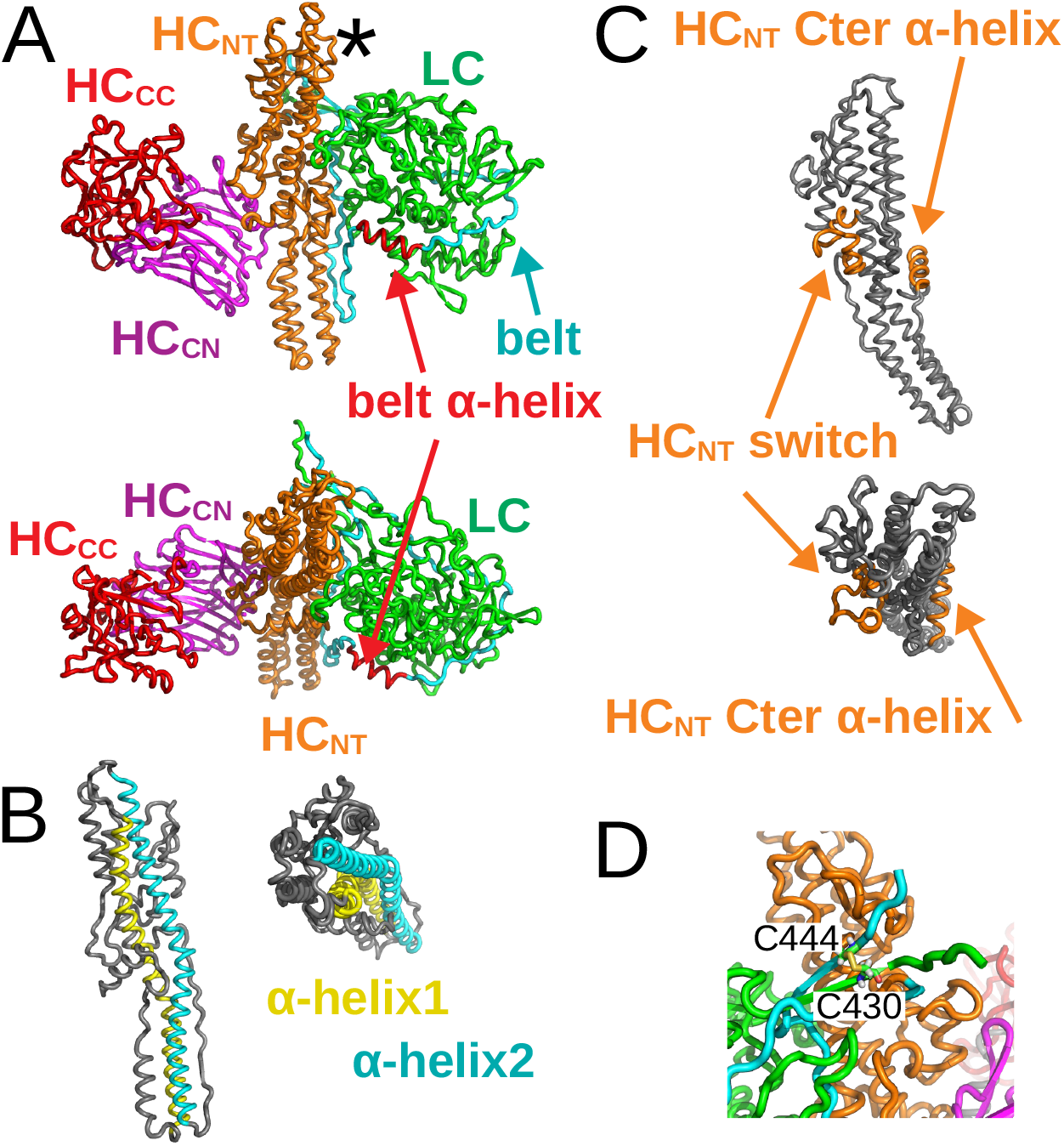
X-ray crystallographic structure of BoNT/A1 in open state drawn in cartoon with various domains in colors (PDB entry: 3BTA [7]). A. Full view of the structure with domains LC (green), belt (cyan), HC_NT_ (orange), HC_CN_ (magenta) and HC_CC_ (red). The belt *α* helix is colored in red. The top of the HC_NT_ domain, close to the disulfide bridge, is indicated with an asterisk. Top: side view. Bottom: upper view. B. Translocation domain in BoNT/A1 structure with the *α* helix1 (cyan) and helix2 (yellow). Left: side view. Right: upper view. C. Translocation domain in BoNT/A1 structure with the HC_NT_ switch and the C-terminal *α* helix in orange. Top: side view. Bottom: upper view. D. Zoom on the disulfide bridge between C^430^ and C^444^ and connecting the two chains.

Furthermore, the X-ray crystallographic structures display two distinct conformations, termed open and closed, characterized by different arrangements of the LC and receptor binding domains with respect to the central HC_NT_ helical domain. In the open conformation, the LC and receptor binding domains are far apart, as open wings lying on either sides of HC_NT_ (see Figure 1A). In the closed conformation, LC and receptor binding domains come in close contact, as closing wings of a butterfly [10]. The open conformation was observed in most of the X-ray structures [7], whereas the closed conformation was only observed for the structure of BoNT/E1 [9]. Some of the X-ray crystallographic structures were determined at acidic pH values in the range 4 to 6 [8], but no structural variation has been observed. Interestingly, BoNTs closely related tetanus toxin TeNT, displays a closed conformation that shows a different organization compared with BoNT/E1 [11,12]. BoNT/A associates with the non-toxic non-hemagglutinin (NTNH) protein at acidic pH to form stable complexes resistant to protease and acidic degradation [13]. Investigation of the BoNT/A-NTNH assembly by small angle X-ray scattering (SAXS) revealed BoNT/A conformational intermediates between open and closed ones [14].

As for their functional mechanism, when approaching the terminal button of target neurons, BoNTs recognize two distinct classes of receptors [15]: complex gangliosides [16], specifically expressed on vertebrate presynaptic neuronal membranes, and synaptic vesicle SV2 for BoNT/A1 and BoNT/E1 [17,18] or synaptotagmin for BoNT/B [19]. BoNT/A1 is able to utilize all three SV2s (A,B,C), whereas BoNT/E1 uses only SV2A or SV2B, but not SV2C, at least not in cultured neurons [18]. Receptor-binding by BoNTs is followed by the endocytosis of the toxin within recycling synaptic vesicles. Following the step of endocytosis, a pH acidification within the vesicle interior triggers toxin-mediated translocation of LC through the endosomal membrane, using a mechanism whose molecular bases have still to be understood [20–22]. Once BoNTs-LC-Zn^2+^-metalloproteases are delivered into the cytosol of neuron presynaptic endings, they cleave a protein of the SNARE complex [23]. As the SNARE complex constitutes the central component of calcium influx-mediated fusion of synaptic vesicles for neurotransmitter release, their cleavage in cholinergic neurons by LC-BoNTs leads to the deadly flaccid paralysis [24].

Back to the comparison between BoNT/A1 and E1, the difference recorded in the onset and duration of the paralysis between these two subtypes covers contrasted behaviors at the level of the various physiological steps underlying the mechanisms of action of these two neurotoxins. Notably, after injection of botulinum neurotoxins BoNT/A and BoNT/E in the muscle of patients, the neuromuscular junction recovers more rapidly from BoNT/E paralytic effect than BoNT/A [5]. It was also reported that BoNT/E LC is degraded more rapidly by the ubiquitin-proteasome system in the cytosol, as compared to BoNT/A LC which is relatively stable [25]. Increased stability of BoNT/A LC involves the activity of the debiquitinating enzymes, VCIP135 that prevents BoNT/A LC degradation by the proteasome and USP9X that prevents its lysosomal degradation [26]. The stabilized form of BoNT/A co-localizes with SNARE components at the presynaptic membrane while BoNT/E LC localizes into the cytosol [27,28]. Moreover, the translocation of the catalytic domain of BoNT/A from the acidified lumen of endosomal compartments to the cytosol was shown to be slow relative to that of BoNT/E [29].

This finding calls for capturing the molecular dynamics of BoNT translocation across endosomal membranes. In this respect, some internal dynamics has been described in recent studies of BoNTs and TeNT. First, a SAXS study of TeNT showed that, at acidic pH, the gyration radius (RG) decreases, an observation which might be related to the appearance of a closed conformation [11]. In addition, a region of HC_NT_, which was named as the HC_NT_-switch, was shown to display conformational transitions enabling membrane insertion of the translocation domain. [30]. Recently, an analysis of BoNT/B and BoNT/E structures by Cryo-electron microscopy (Cryo-EM) [31] showed that these structures are more mobile than the corresponding X-ray crystallographic structures. Although some of these structures might be not all functional, these observations point toward an internal mobility of BoNTs, compared with the relative structure uniformity of BoNTs recorded by X-ray crystallography.

Molecular level information on toxins internal mobility can be obtained by atomistic simulations [32], which could provide insight on the different possible conformations of BoNTs, and on how these are affected by the environment. Few molecular dynamics (MD) simulations of BoNTs have been reported. A study has been reported on the full-length BoNT/A in water at different pH and temperature [33] while in another study the interaction of the BoNT/A receptor binding domain with the synaptic vesicle protein 2C (SV2) luminal domain was investigated [34]. A recent study [35] focused on the pH-dependent structural changes in BoNT/E1, based on extensive MD simulations at various pH values, and on small angle X-ray scattering analysis.

The present work intended to explore the internal dynamics of two full-length BoNTs in a large water system and along hundred of nanosecond time scale: BoNT/A1 and BoNT/E1, using the information provided by X-ray crystallographic structures along with homology modeling. Starting conformations of BoNT/A1 and E1 were built to analyze the behavior of open and closed states of the toxins, at neutral and acidic pH conditions. The aim is at investigating the protein internal dynamics at the ternary and quaternary levels, and to relate the observed conformational changes to relevant physiological steps. A general mechanism is proposed for the initiation of translocation in BoNTs. A comparative analysis of the cleaved toxins BoNT/A1 and E1 provides a first-level description of putative structural determinants for the different translocation kinetics driven by intraluminal acidification in these two BoNT serotypes.

## 2. Results

In this section, we present the results obtained by analyzing several descriptors of protein’s structure and dynamics along the MD trajectories (Table 1) recorded starting from cleaved X-ray crystallographic conformations or from trans models, as described in detail in section 4.1. To anticipate the meaning, the term ’trans models’ here refers to structures fully obtained by homology modeling calculations (e.g. open conformations of E1 and closed conformations of A1), to be distinguished from conformation modeled on the available experimental X-ray structures (e.g. closed E1 and open A1 conformations, respectively.)

**Table 1.** Systems composition for molecular dynamics simulations. The names of the systems, reported in the first column, are introduced in Section 4.1. “Preparation of starting conformations of toxins”. The characters “clo” and “ope” refer to the closed and open states of BoNTs, the characters “47” and “70” refer to the pH values and the character “r” refers to the trans models determined under distance restraints of Table S3.

| System | number of water molecules | counter-ions | total number of atoms |
| --- | --- | --- | --- |
| A1clo47 | 162483 | 23 Cl<sup>-</sup> | 508335 |
| A1clo47r | 162409 | 28 Cl<sup>-</sup> | 508123 |
| A1clo70 | 162514 | 2 Cl<sup>-</sup> | 508386 |
| A1clo70r | 162418 | 5 Cl<sup>-</sup> | 508104 |
| A1ope47 | 162499 | 19 Cl<sup>-</sup> | 508373 |
| A1ope70 | 162531 | no counter-ions | 508431 |
| E1clo47 | 162749 | 24 Cl<sup>-</sup> | 508408 |
| E1clo70 | 162777 | 2 Cl<sup>-</sup> | 508448 |
| E1ope47 | 162634 | 35 Cl<sup>-</sup> | 508083 |
| E1ope47r | 162639 | 32 Cl<sup>-</sup> | 508092 |
| E1ope70 | 162673 | 3 Cl<sup>-</sup> | 508136 |
| E1ope70r | 162668 | 6 Cl<sup>-</sup> | 508127 |

We started with the analysis of residues protonation, which reflects both the pH and the peculiar BoNT conformation (open or closed). Residue protonation, defined at the modeling stage, has direct effect on the protein internal dynamics, determining intra- and inter-domain long-range interactions. The Root Mean Square Deviation (RMSD) between conformations as well as the distances between domains were utilized to get global information on structural rearrangements in BoNTs with time. Atomic Root Mean Square Fluctuations (RMSFs) were analyzed to define local motions in distinct domains, while monitoring inter and intra-domains hydrogen bonds provides information complementing the global structural results. We then focused on the dynamics of three domains: the belt, the core translocation domain and the binding receptor domains.

Definition of BoNT domains utilized in this work is reported in Table 2.

**Table 2.**
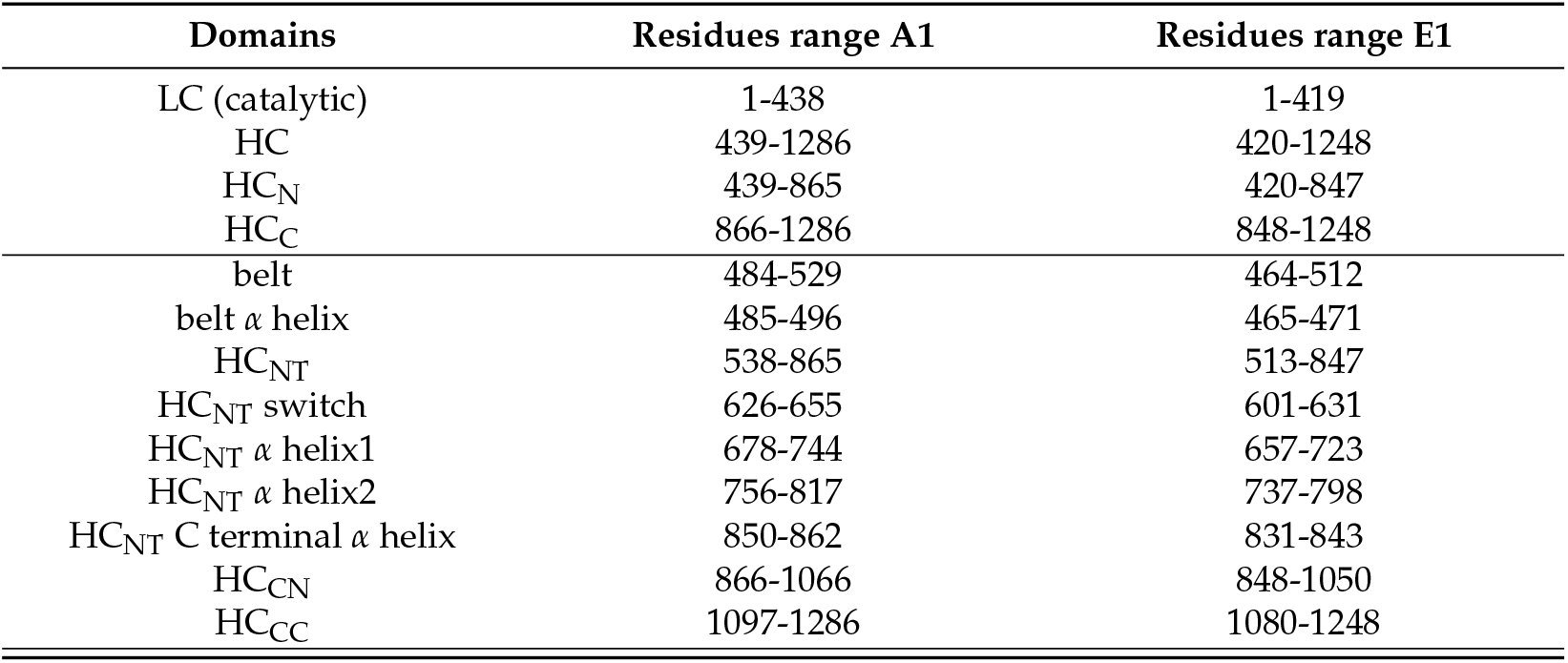
Definition of domains used here for BoNTs A1 and E1.

| Domains | Residues range A1 | Residues range E1 |
| --- | --- | --- |
| LC (catalytic) | 1-438 | 1-419 |
| HC | 439-1286 | 420-1248 |
| HC<sub>N</sub> | 439-865 | 420-847 |
| HC<sub>C</sub> | 866-1286 | 848-1248 |
| belt | 484-529 | 464-512 |
| belt $\alpha$ helix | 485-496 | 465-471 |
| HC<sub>NT</sub> | 538-865 | 513-847 |
| HC<sub>NT</sub> switch | 626-655 | 601-631 |
| HC<sub>NT</sub> $\alpha$ helix1 | 678-744 | 657-723 |
| HC<sub>NT</sub> $\alpha$ helix2 | 756-817 | 737-798 |
| HC<sub>NT</sub> C terminal $\alpha$ helix | 850-862 | 831-843 |
| HC<sub>CN</sub> | 866-1066 | 848-1050 |
| HC<sub>CC</sub> | 1097-1286 | 1080-1248 |

### 2.1. Analysis of the protonation at varying pH in different states

The protonations of amino-acid residues for the various states and at different values of pH are listed in Tables S1 and S2. In addition to the three protonation states of histidines, protonated histidines on N*δ* (HSD), protonated histidines on N*ϵ* (HSE), doubly protonated histidines (HSP), the other protonated residues are: glutamate (GLU) and the aspartate (ASP), in which an hydrogen was added to the sidechain carboxyl group.

Different numbers of protonated residues are observed in the BoNT domains. A large number of protonated residues, in the range 7-13 for BoNT/A1 and in the range 6-12 for BoNT/E1, is observed in LC. The number of protonated residues increases from around 8 at neutral pH to the range 10-13 at acidic pH, and in this condition, it is larger in the closed conformations of BoNT/A1 (around 13) than in the open ones. Another large cluster of protonated residues is located at acidic pH in the domain HC_NT_, in the range 9-14, and, as for LC, this number increases with acidic pH and is larger in the closed conformations. Some of these residues are located in the HC_NT_ switch (D^640^, E^656^ in A1clo47r and E^616^, E^623^, E^632^ in E1ope47). Others (D^838^ in A1clo47r, D^829^ in E1clo47 and E1ope47) are close to the C terminal helix of HC_NT_. In BoNT/E1, other residues are located close to the *α* helix1 and helix2, in particular to residue K^775^, E^636^, E^677^ in E1clo47 and E^773^, E^781^ in E1ope47 and E1ope47r. Overall, only few residues are protonated in the belt and in the domains HC_CN_ and HC_CC_, except for the open state of BoNT/E1 at acidic pH (E1ope47 and E1ope47r).

### 2.2. Variations of the intra- and inter-domain organization in BoNTs

The combined analysis of RMSD, RMSFs and hydrogen bond networks provides useful insights for tertiary and quaternary modifications in different conformational and protonation states.

The RMSD of C*α* atoms along the MD trajectories (Figure 2) show plateaus in some cases already after 50 ns. Plateau values are up to 7 Å for the whole structure (top plots), with a temporary jump up to 10 Å around 125 ns in the trajectory E1clo70 (green curve).

**Figure 2.**
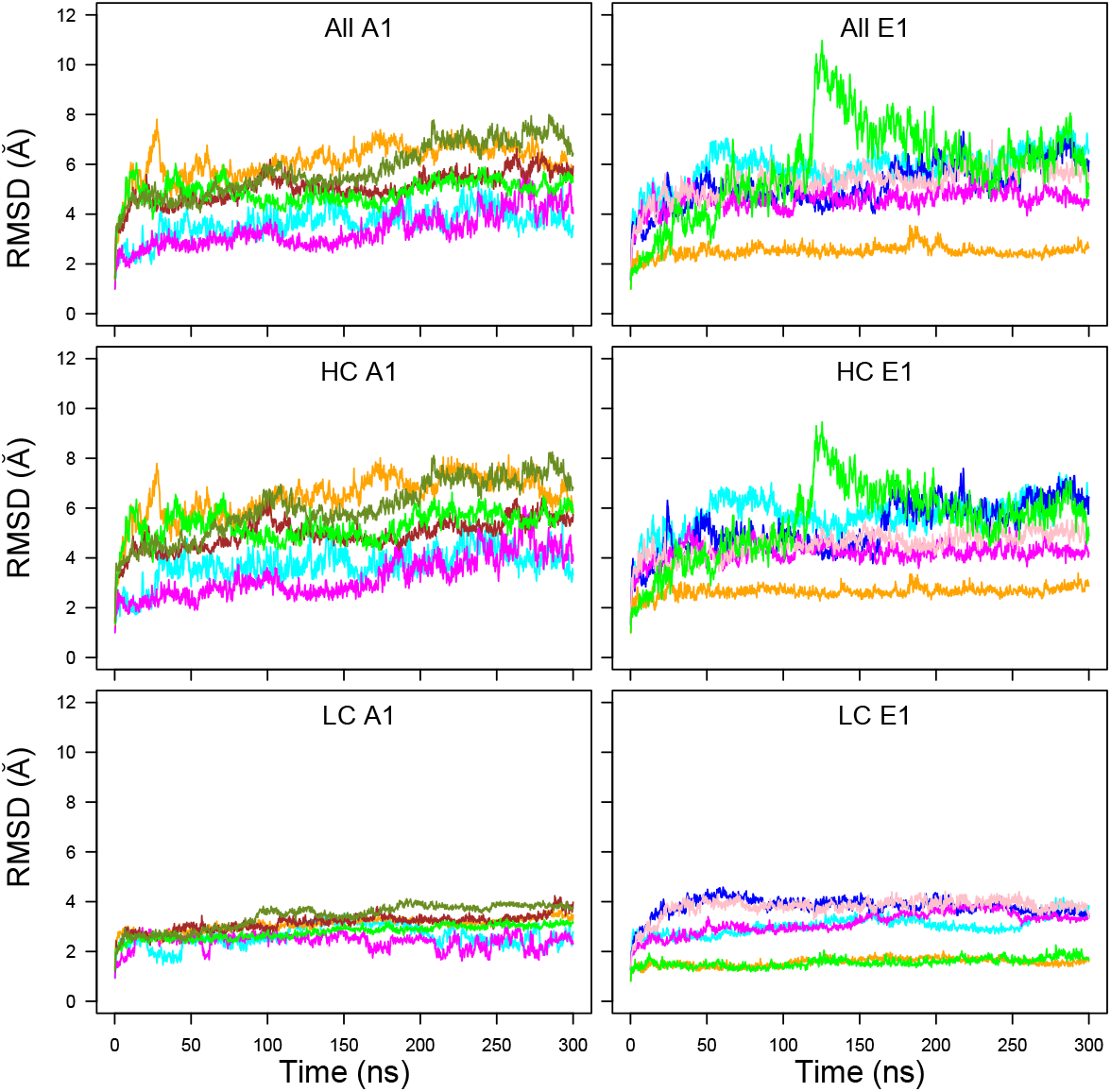
C*α* root-mean-square deviation (RMSD, Å) along the MD trajectories recorded on the botulinium toxins A1 (left column) and E1 (right column). The curves are plotted with colors: cyan (A1ope47, E1ope47), blue (E1ope47r), magenta (A1ope70,E1ope70), pink (E1ope70r), orange (A1clo47, E1clo47), brown (A1clo47r), green (A1clo70, E1clo70), olive green (A1clo70r). The RMSD are plotted for the whole structure (top row), the heavy chain (HC, middle row) and the light chain (LC, bottom row).

Very flat and low profiles around 2-4 Å are observed for the light chain (LC, bottom plots) whereas the RMSD calculated on the heavy chain (HC, middle plots) dominates the total RMSD values. The RMSD values for LC are in the range 2-4 Å for A1, whereas for E1, some RMSD curves are smaller than 2 Å, the others being in the range 3-4 Å. The LC, composed mostly of the catalytic domain, displays thus a stable conformation.

The BoNT/A1 trajectories starting from a trans model display RMSD values in the 4-6 Å range (green, orange, olive green, and brown curves). These values are slightly larger than those observed for the open state trajectories generated from X-ray cleaved models (magenta and cyan curves). A similar feature is observed for the BoNT/E1, with a larger gap between cleaved X-ray (orange and green curves) and trans models (blue, pink, magenta and cyan curves). In that case, the use of additional restraints, described in section 4.1 and in Table S3, to enforce the interaction between the belt *α* helix and its environment (pink and blue curves) does not actually reduce the RMSD.

The trajectories starting from X-ray cleaved models (A1ope47, A1ope70, E1clo47, E1clo70) display different behaviors in A1 and E1. Indeed, unlike BoNT/A1, the trajectory E1clo70 (green curve) displays a large jump around 125 ns, in which the receptor-binding domains (HC_CN_ and HC_CC_) and the LC domain move slightly apart (Figure S1). This lack of stability could arise from a bias in the X-ray crystallographic structure (3FFZ) induced by the crystal packing or by the use of a unique protein chain to produce the sample for crystallographic purposes.

Nevertheless, protonation effects could possibly play a role. Indeed, in analogy with the conformational transition observed experimentally at acidic pH value of 5.0 for TeNT [11], one could conceive that the open form of A1 at pH 7.0 (magenta curve) would be more stable than at pH 4.7 (cyan curve), while on the contrary the closed form of E1 would be more stable at pH 4.7 (orange curve) than at pH 7.0 (green curve). Actually, it is what is observed for the HC RMSD in Figure 2, by comparing the A1 magenta and cyan curves (A1ope70 vs A1ope47), and the E1 orange and green curves (E1clo47 vs E1clo70), respectively. To resume, two factors would contribute to the protein internal stability: artifacts of the homology modeling templates, plus protonation effects which shift the equilibrium toward one or the other conformation of the protein.

The distributions of C*α* RMSD calculated for the individual BoNT domains are displayed as boxplots (Figure 3). We found that the RMSD values, except for belt and HC_CC_, cluster in a much narrower range (1-4 Å for the medium values) than the global RMSD values observed in Figure 2. This observation supports a model of mobility in which the individual domains fluctuate around stable conformations and most of the motions of the overall structure arise from relative displacements of the domains. This agrees with previous results from molecular dynamics simulations on BoNTs [33,35]. In this frame, the outlier values observed for the belt in the closed state of BoNT/A1 are not surprising as this region is an extended loop connecting the catalytic and the HC_NT_ domains. One should also notice that HC_CC_ explores much larger RMSD values than the other domains, specially for the closed state of BoNT/A1. LC, HC_NT_ and HC_CN_ display smaller RMSD values.

**Figure 3.**
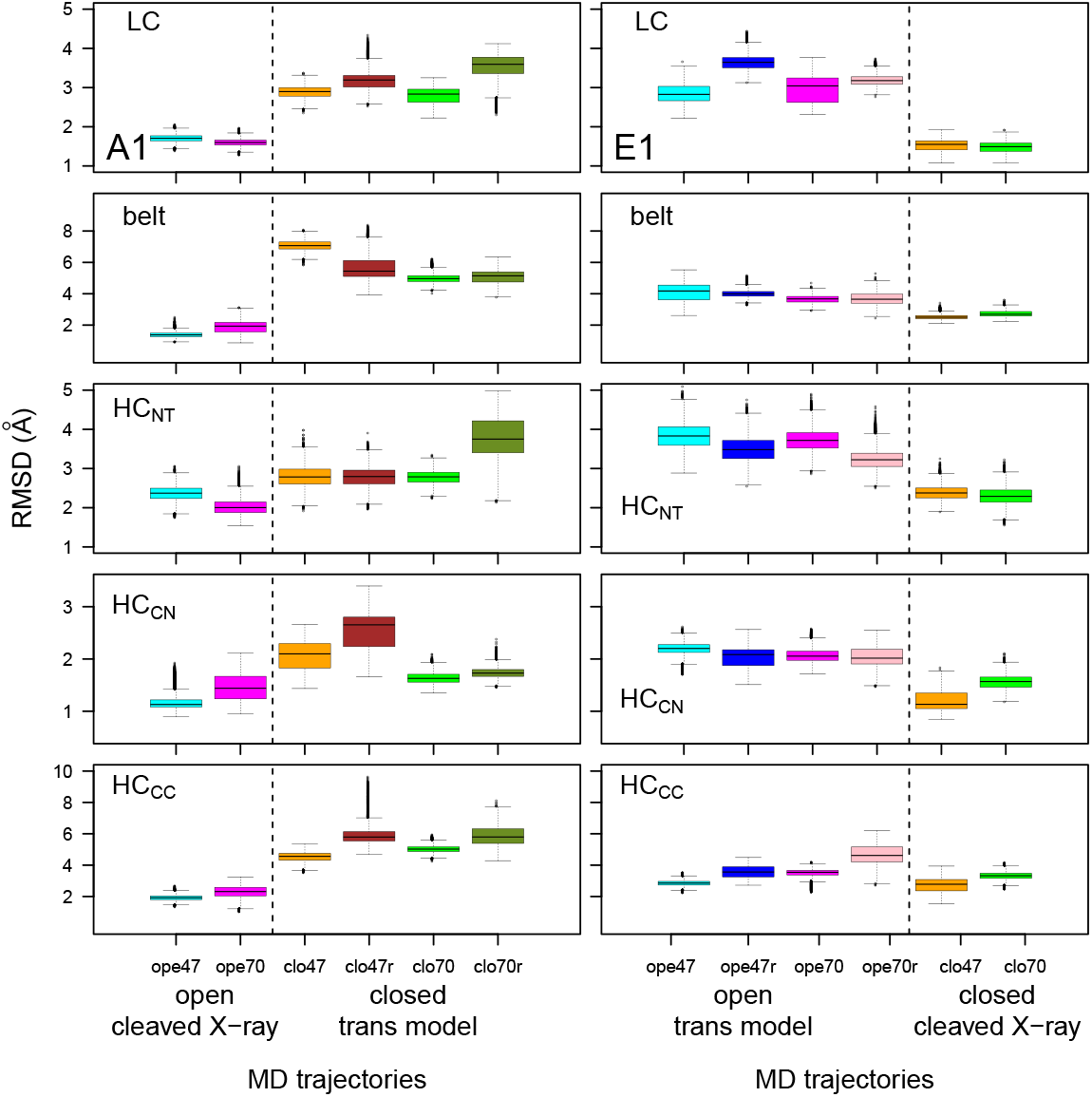
Box-and-whisker plot representation of the distributions of RMSD (Å) for domains of the botulinum toxins A1 and E1. The color code of the boxes is the following: cyan (A1ope47, E1ope47), blue (E1ope47r), magenta (A1ope70,E1ope70), pink (E1ope70r), orange (A1clo47, E1clo47), brown (A1clo47r), green (A1clo70, E1clo70), olive green (A1clo70r). The dashed lines mark the separation between open and closed states, thus allowing to show results relative to the various structures in the same graph, providing in this way an overall view of the changes induced by the closing/opening, by the inclusion of restraints in modeling as well as by pH.

A repeated feature in Figure 3 is the increase of RMSD values for MD trajectories starting from trans models with respect to MD trajectories starting from cleaved X-ray models. This has been already observed for global RMSD (Figure 2) and is related to the percentage of identity in the 35-45% range between the primary sequences of the two toxins. Some exceptions are nevertheless observed, as for the belt and HC_CC_ displaying similar RMSD values in all E1 trajectories.

RMSFs along the LC and HC residues (Figure 4) are analogous for BoNT/A1 and BoNT/E1 with peaks observed at similar positions. Two peaks are observed at the two extremities of the domain HC_NT_ for residues 746-751 (A1) and 723-734 (E1) (peak a) located in the loop at the top of HC_NT_ indicated by asterisk in Figure 1A, and for residues 813-826 (A1) and 798-805 (E1) (peak b) located in the loop at the bottom of HC_NT_. Another peak is observed in HC_NT_ for the residues 632-656 (A1) and 601-656 (E1) (peak c) located in the HC_NT_ switch and in the linker connecting the HC_NT_ switch and the *α* helix2. The domain HC_CC_ displays numerous peaks of mobility for both toxins, the largest is observed for A1clo70r and indicated with the letter d.

**Figure 4.**
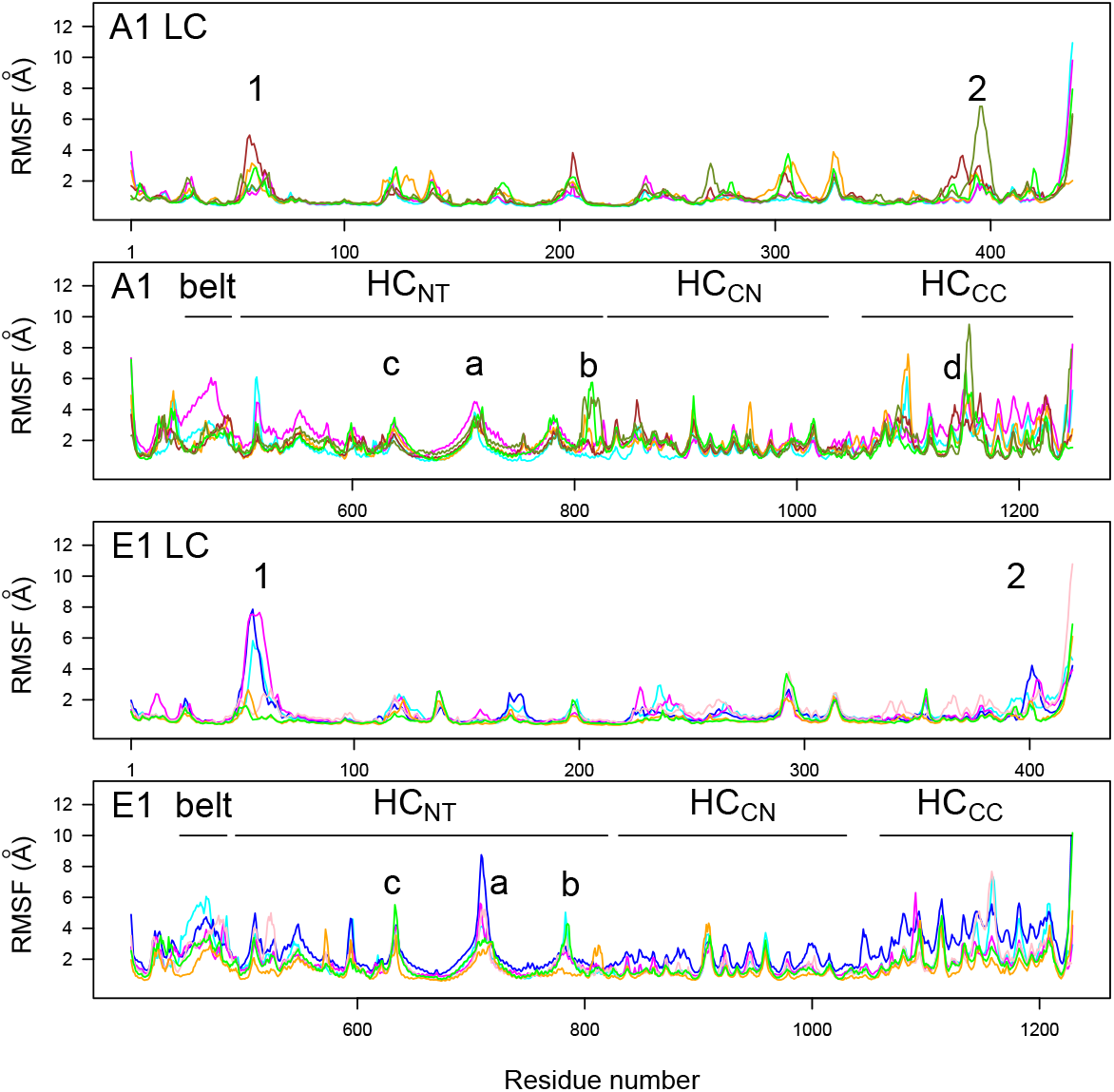
C*α* root-mean-square fluctuations (RMSF, Å) calculated on the interval 150-300 ns of MD trajectories recorded on the botulinum toxins A1 and E1. The various domains (LC, HC_NT_, HC_CN_ and HC_CC_) are indicated on the plots. The color code of the curves is the following: cyan (A1ope47, E1ope47), blue (E1ope47r), magenta (A1ope70,E1ope70), pink (E1ope70r), orange (A1clo47, E1clo47), brown (A1clo47n), green (A1clo70, E1clo70), olive green (A1clo70r). Peak 1: residues 63-65 (A1) and 53-60 (E1); peak 2: residues 393-394 (A1) and 392-399 (E1). Peak a: residues 746-751 (A1) and 723-734 (E1); peak b: residues 813-826 (A1) and 798-805 (E1); peak c: residues 632-656 (A1) and 601-656 (E1); peak d: residue 1188-1198 (A1).

LC displays only two peaks in the RMSF profiles, with values larger than 3 Å. The peak 1 is located around the residues 63-65 for A1 and 53-60 for E1 and corresponds to a loop on the top of the catalytic domain. The peak 2 is located around the residues 393-394 for A1 and 392-399 for E1 and corresponds to a region just before the disulfide bridge.

Distances between geometric centers of several domains have been monitored (Figure 5) along MD trajectories. Distances between LC and HC_NT_ domains and between HC_CN_ and HC_CC_ display similar values across closed and open forms, cleaved X-ray models or trans models, and along pH changes. On the other hand, the distances between HC_NT_ and HC_CN_ or HC_CC_ domains increase when comparing the open and closed conformations, regardless of whether the starting point is a cleaved X-ray model or a trans model. This increase is nevertheless different between the two toxins: BoNT/A1 displays a continuous increase whereas BoNT/E1 displays a jump. The moving apart of HC_CN_ and HC_CC_ from the translocation domain might have a functional meaning. Indeed, BoNTs initially interact with receptors through the HC_CC_ domain for BoNT/A1 and through the HC_CN_ and HC_CC_ domains for BoNT/E1 [36], and then translocate through the vesicle endosomal membrane. Moving apart HC_CN_ and HC_CC_ from the HC_NT_ domain could free HC_NT_ and catalytic domains and make them available for the translocation through the vesicle membrane. In this picture, the closed state induced by acidic pH [11] would be the most prone to translocate. In addition, by assuming that the increase of the HC_NT_/HC_CN_ and HC_NT_/HC_CC_ distances is an early event required for translocation, the steepest distance jump observed in E1 could support at molecular level the experimental observation that pH-dependent translocation of BoNT/E is fast relative to that of BoNT/A [29].

**Figure 5.**
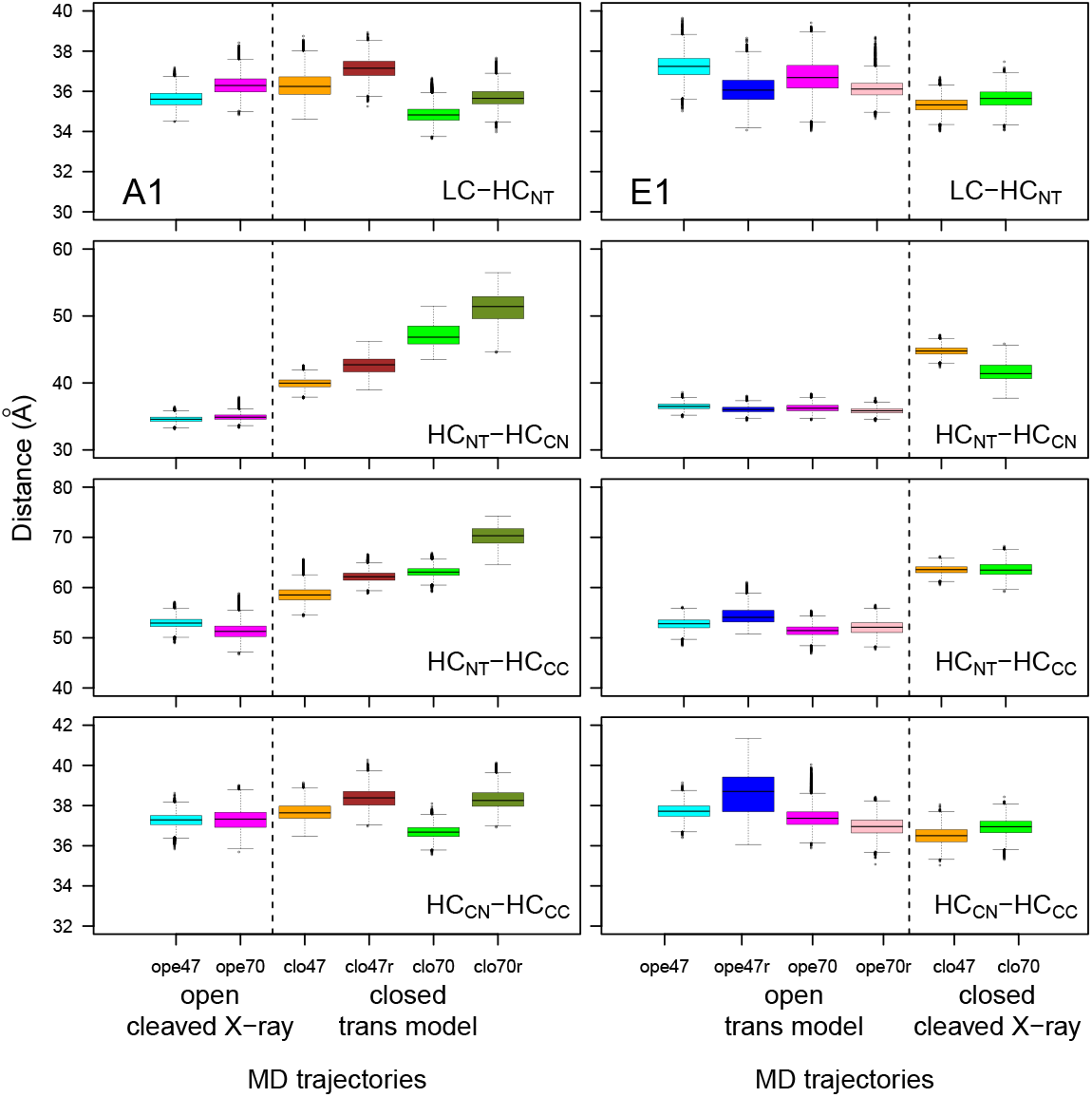
Box-and-whisker plot representation of the distributions of distances between the geometric centers of the domains within the botulinum toxins A1 and E1, averaged on the interval 150-300ns of the MD trajectories. The color code of the boxes is the following: cyan (A1ope47, E1ope47), blue (E1ope47r), magenta (A1ope70,E1ope70), pink (E1ope70r), orange (A1clo47, E1clo47), brown (A1clo47r), green (A1clo70, E1clo70), olive green (A1clo70r). The dashed lines mark the separation between open and closed states, see Figure 3.

Long-range hydrogen bonds were detected along the trajectories using the python package MD Analysis [37,38]. The number of hydrogen bonds present more than 60% of the time and involving residues separated by more than 10 residues in the sequence was calculated between and within BoNT domains (Figure 6). The number of long-range hydrogen bonds within the LC domain, and to a lesser extent within HC_CN_, is high in all trajectories (displaying however a slightly decrease in trans models), and this agrees with the smaller RMSD values observed for LC and HC_CN_ (Figure 3). By contrast, the number of hydrogen bonds between the belt and LC domains and within HC_NT_ and HC_CC_ is mostly smaller and decreases for trajectories starting from trans models. The smaller number of hydrogen bonds within HC_CC_ agrees with the large RMSD (Figure 3) and RMSF (Figure 4) values of this domain.

**Figure 6.**
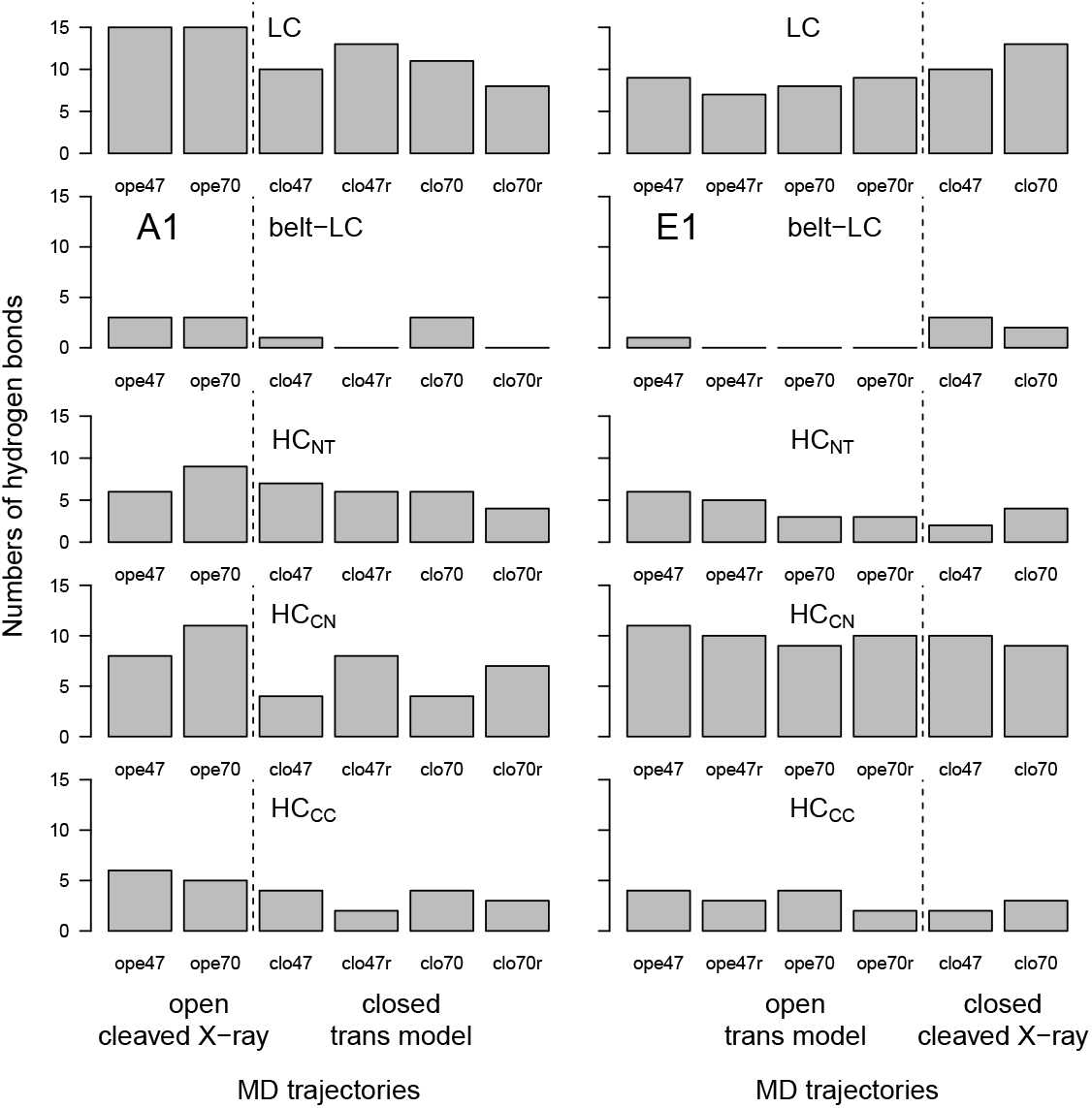
Number of hydrogen bonds between domains of the botulinum toxins A1 and E1 calculated on the interval 150-300 ns of trajectories. Long-range hydrogen bonds were detected along the trajectories by analyzing each frame every 10, using the python package MDAnalysis [37,38]. The numbers of hydrogen bonds are calculated as those present more than 60% of the time and involving residues separated by more than 10 residues in the sequence. The dashed lines mark the separation between open and closed states, see Figure 3.

### 2.3. Flexibility of the lipid-binding and ganglioside-binding domains in HC_CC_

The analysis of the root-mean-square fluctuation profiles revealed large internal mobility in the domain HC_CC_ (Figure 4). In particular, a peak, labeled d, is observed for loop 1188-1198 in the domain HC_CC_ for the trajectory A1clo70r, much larger than any such peak observed for E1. In the initial conformation of HC_CC_, the loop 1188-1198, which in A1 corresponds to the lipid-binding loop (LBL), adopts a *β* strand conformation, forming a small *β* sheet with the *β* strand spanning residues 1252 to 1254 (sequence VAS) close and partially overlapping to the stretch of residues 1254-1257 (SNWY). Similarly, the corresponding loop in E1 (residues 1164-1179) establishes in the initial conformation of HC_CC_ a *β* sheet with the stretch of residues VAS (residues 1216-1218), close to the stretch of residues STWYY (1218-1222). Remarkably, the stretch SXWY is the conserved ganglioside binding site (GBS) and has been identified first in tetanus neurotoxin as well as in BoNT/A, B, E, F, and G [39–41]. Interestingly, in E1 GBS, key interacting residues that are unique to BoNT/E have been identified along with a significant rearrangement of loop 1228-1237 upon carbohydrate binding [42]. In BoNT/B, the LBL is located between the GBS and the receptor binding site, and is significantly exposed to the solvent, in particular the sidechains of W^1248^ and W^1249^ [43]. In the X-ray crystallographic structures of BoNTs, the regions LBL and GBS are close in the 3D space and correspond to well defined binding regions (see e.g. in Figure S2 for A1 in the open state (PDB entry: 3BTA).

The distributions of distances corresponding to backbone hydrogen bonds between the two *β* strands initially present in the X-ray crystallographic structures were monitored along MD trajectories (Figure 7). In the open state of A1, the interactions between *β* strands 1188-1198 and 1252-1254 established in the starting point are kept during the trajectory (see Figure S2). In all other trajectories, the interaction between *β* strands is lost. In general, the mobility observed in this region during simulations is more pronounced in the case when additional restraints (Table S3) are applied on the *α* helix, specially for the closed state in A1 (A1clo70r, olive green box) and for the open state of E1 (E1ope70r, pink box).

**Figure 7.**
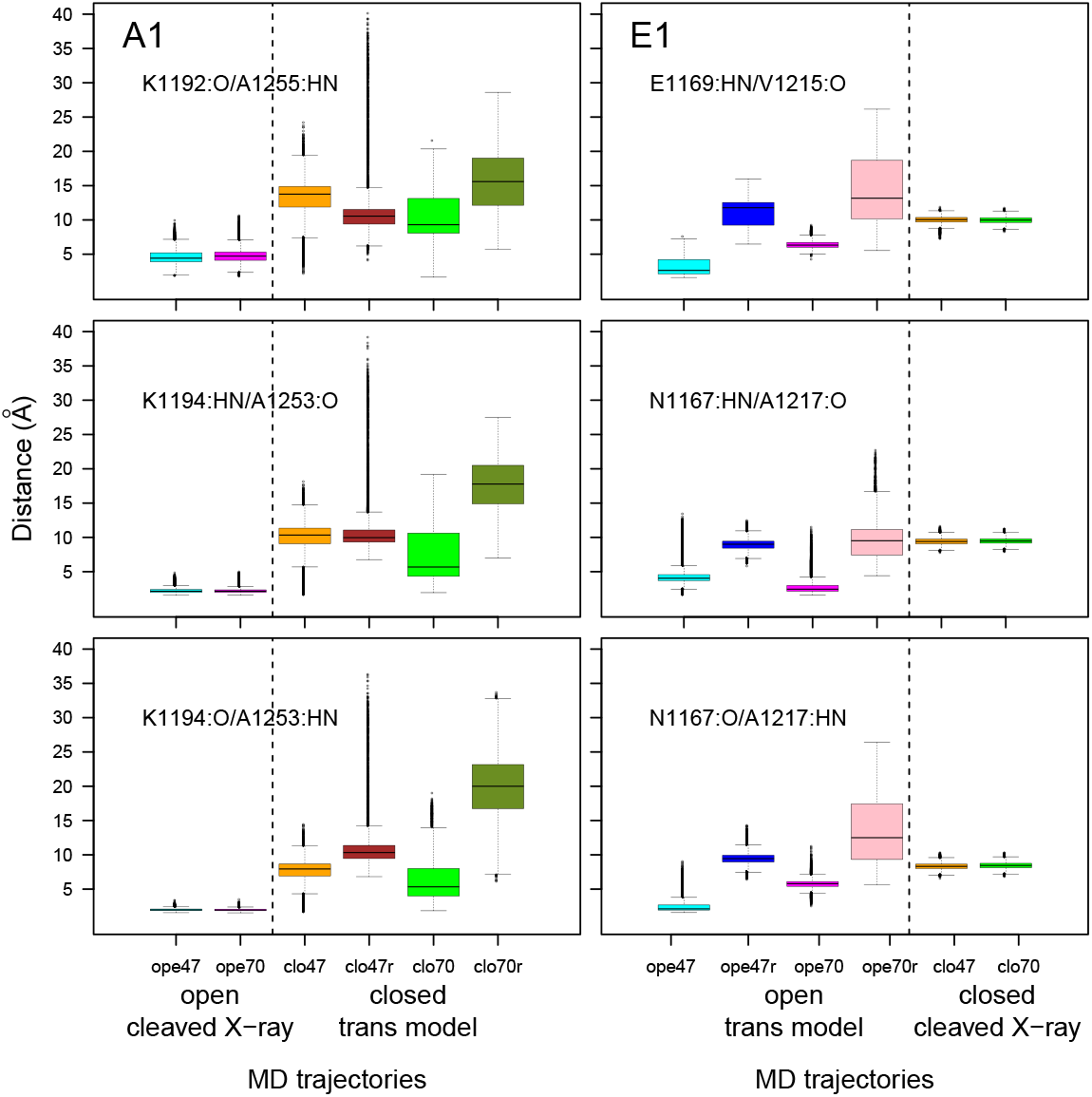
Box-and-whisker plots representation of distributions of distances corresponding to the hydrogen bonds between the amide hydrogens (HN) and carbonyl oxygens (O) of residues 1188-1198 (LBL) and 1252-1254 (close to the GBS) for A1 and of residues 1164-1179 (LBL) and 1216-1218 (close to the GBS) for E1. The color code of the boxes is the following: cyan (A1ope47, E1ope47), blue (E1ope47r), magenta (A1ope70,E1ope70), pink (E1ope70r), orange (A1clo47, E1clo47), brown (A1clo47r), green (A1clo70, E1clo70), olive green (A1clo70r). The dashed lines mark the separation between open and closed states, see Figure 3.

The mobility of the loop LBL in BoNT/A1 and E1 may look paradoxical, as the X-ray crystallographic structures determined so far were showing quite well defined interaction pockets with protein receptor and gangliosides [44]. Overall, the very mobile domain HC_CC_ observed here points to a different interaction mechanism than the one inferred from the X-ray crystallographic structures. Indeed, a recent closer investigation of the interaction between BoNT/B, ganglioside GT1b and synaptotagmin has revealed [44] that a complex GT1b-synaptotagmin preexists to the BoNT/B binding, stabilizing the conformation of the synaptotagmin juxtracellular domain, in a way quite different to what was observed in the structures. The high flexibility of the HC_CC_ domain observed in our models is consistent with this variability of BoNT conformations and of their interaction with receptors. It should be nevertheless noticed that the present study did not investigate the binding of BoNTs with the N glycans.

### 2.4. Conformational variations in the core translocation and belt domains

To investigate the mobility of the core translocation and belt domains, several structural descriptors have been analyzed whose combination would help in dissecting possible earliest events of the translocation. In particular we analyzed the solvent accessible residue surfaces, the behaviour of the backbone dihedrals in the belt domain and in the HC_NT_ switch region, as well as the bending of the *α* helices 1 and 2 in HC_NT_.

#### 2.4.1. Solvent accessible residue surface

The solvent accessible surfaces of residues were calculated along the trajectories and the time-averaged values clustered according criteria detailed in Section Material and Methods. We could group the residues in four sets: i) residues more accessible in closed than in open state, at both pH; ii) residues more accessible in open state than in closed state, at both pH; iii) residues more accessible in closed state at pH 7.0 than in other states; iv) residues more accessible in closed state at pH 4.7 than in other states. Residues belonging to the four sets are listed in Table 3. Their position along the protein sequence is shown in Figure S3. To convey the information where patches of more exposed residues are located, in the different subtypes and pH conditions, residues are displayed in Figure 8 on the structures of the open conformations of BoNT/A1 and BoNT/E1.

**Table 3.** Residues displaying variations of accessible surfaces between the different systems. Protonated residues quoted in Tables S1 and S2 are written in bold.

|  |  |
| --- | --- |
| System A1 | residues more accessible in closed than in open state<br>LC: M30 R48 T52 I154 N205 G255 N353 I376 F419 T420<br>HC: L465<br>HC<sub>NT</sub>: N560 K616 A618 D619 G644 F648 V783 N788 I791<br>K796 **E799** N816 L843 I863<br>HC<sub>CN</sub>: L918 |
| System A1 | residues more accessible in closed state at pH 7.0 than in other states<br>LC: I45 G119 G120 Y180 I348 F357 N362 N402<br>HC: N449<br>HC<sub>NT</sub>: I737<br>HC<sub>CC</sub>: M1124 V1176 |
| System A1 | residues more accessible in closed state at pH 4.7 than in other states<br>LC: F36 I111 P116 P156 **H170** G179 T183 A228 L232 Y233 Y251 F290<br>K320<br>belt: F498 G516<br>HC<sub>NT</sub>: R692 L809<br>HC<sub>CN</sub>: I922 V923 Y924 N944<br>HC<sub>CC</sub>: T1135 F1150 I1172 G1220 G1238 G1241 F1252 |
| System A1 | residues more accessible in open state than in closed state<br>LC: N5 **H39** A65 Y72 G211 I237 K375 L422<br>belt: Q491 L494 N500 N509<br>HC<sub>NT</sub>: I556 L558 T559 V562 V572 T614 K710 A732 I752 I821<br>HC<sub>CN</sub>: L869 S886<br>HC<sub>CC</sub>: R1121 R1169 N1247 I1248 G1269 |
| System E1 | residues more accessible in closed state than in open state<br>LC: T234 Q237 S292 L391 R394<br>belt: I489<br>HC<sub>NT</sub>: S536 L624 A667 W686 I737 **E741** T760 E761 S763 I764<br>K769 N772 **E773** K775 I776 N777 I794 I800 N809 V812 L824 T828 L833<br>HC<sub>CN</sub>: **D866** K908 N985<br>HC<sub>CC</sub>: F1157 |
| System E1 | residues more accessible in open state than in closed state<br>LC: L144 T228 Q235 N392 I396 V405<br>HC: E462<br>belt: T503<br>HC<sub>NT</sub>: V532 I538 K728 P821 K823 N838 K839 F841 K842<br>HC<sub>CN</sub>: S867 N911 F1044<br>HC<sub>CC</sub>: I1172 |

**Figure 8.**
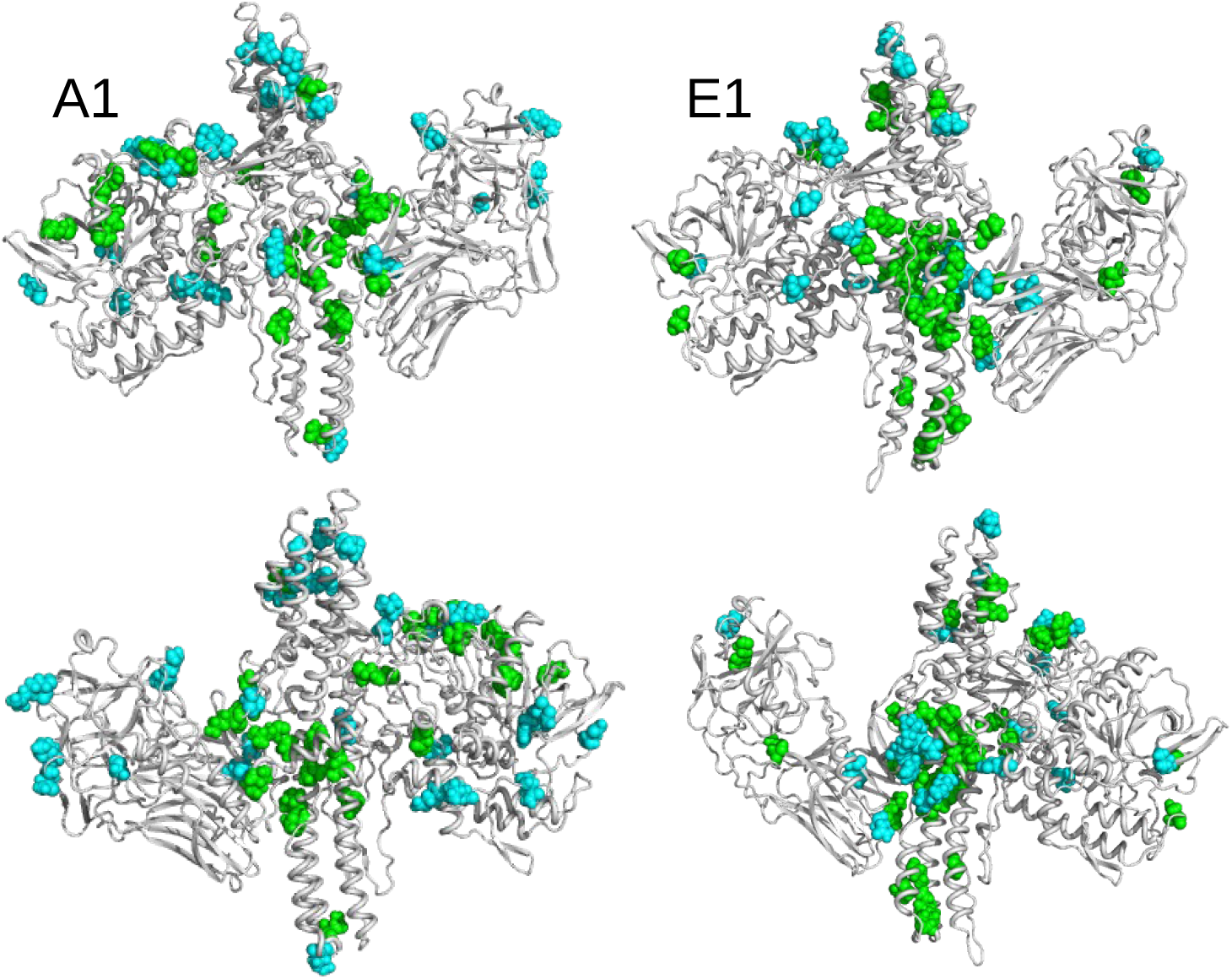
The open conformations of BoNT/A1 and BoNT/E1 are drawn in cartoon and displayed in two opposite orientations. The residues displaying variations of accessible surfaces are shown in van der Waals representation and colored according to the following rules: green residues: more accessible in closed than in open state; cyan: more accessible in open than in closed state.

Overall, for both BoNTs, clusters of green residues more accessible in closed than in open conformations are observed in the translocation domain (HC_NT_) and to a lesser extent in the catalytic domain (LC). On the contrary, the residues colored in cyan, that are more accessible in the open than in the closed conformation, are mostly scattered along the sequence or in the 3D structure.

In details, residues more accessible in the closed state are located in specific regions of HC_NT_, as shown in Figure S4. In BoNT/A1, L^843^ and I^863^ (Table 3) are located at the two extremities of the C terminal *α* helix connecting HC_NT_ and HC_CN_ (Figure 1C), the residue V^783^ is behind the HC_NT_ switch, the residues K^796^ and E^799^ are in the *α* helix2 toward the top of HC_NT_. In BoNT/E1, L^624^ is located in the HC_NT_ switch, E^761^, I^764^ and S^763^ in the helix2, W^686^ in the helix1 and the residues K^769^, N^772^, E^773^, K^775^, I^776^ and N^777^ are located toward the bottom of *α* helix2. These accessible residues observed in the domain HC_NT_ can be related to experimental observations [45] showing that in BoNT/A LC-HCT, residues located in an *α* helix close to the bottom extremity of HC_NT_, display increase in fluorescence intensity and blue shift to 530 nm when I830C-NBD binds to liposomes, in agreement with the transfer into a non-polar environment. Remarkably, among all systems investigated, the number of solvent exposed residues in the HC_NT_ domain is the largest in the closed state of E1. These results would support the hypothesis by Kumaran et al [9] on the faster translocation of BoNT/E [29], which, according to the authors, could be related to the exposure to solvent of one side of the translocation domain.

Among the residues displaying changes in accessible surface (Table 3), several residues change protonation states (Tables S1 and S2) along the pH: H^39^, H^170^ (LC), E^799^ (HC_NT_) and D^866^ (HC_CN_) in A1 and E^741^, E^773^ (HC_NT_) and D^866^ (HC_CN_) in BoNT/E1. In the LC domain, some residues protonated at acidic pH are located in the neighborhood of several residues changing accessible surfaces (Figure S5). Indeed, in BoNT/A1, the residues R^48^, T^52^, I^154^ and N^205^, more accessible in closed than in open state, are respectively in the neighborhood of residues H^39^, D^58^, E^164^ and D^513^ protonated in closed state at acidic pH. In BoNT/E1, the residue S^292^, L^391^ and R^394^, more accessible in closed state, are close in 3D space to H^125^ and E^189^, protonated at acidic pH. As larger accessibility as well as the protonation may facilitate interaction with membrane, their occurrences in residues close in 3D space could also induce a cooperativity in the interaction.

In BoNT/A1, several residues of LC are more accessible in the closed state at acidic pH, namely F^36^, I^111^, P^116^, P^156^, H^170^, G^179^, T^183^, A^228^, L^232^, Y^233^, Y^251^, F^290^ and K^320^. Given that most of them are hydrophobic residues, their exposure to solvent could be related to loosening of the LC domain fold (Figure S6). This could also affect the active site, as A^228^, L^232^, Y^233^ are close to the residues H^223^ and H^227^ of the catalytic site, belonging to the same helix spanning residues 217-233. Such disruption of the LC tertiary structure is also suggested by the high value of the LC RMSD in the closed form at acidic pH (Figure 3, upper left panel) and by the reduction of the intra-protein hydrogen bonds with respect to the open form (Figure 6, upper left panel). The loosening of the LC fold was observed experimentally at acidic pH [46], and could be the first step to prepare its subsequent translocation through the vesicle membrane.

#### 2.4.2. Internal mobility of belt *α* helix and HC_NT_ switch

In order to examine the variations of conformations in the belt domain and HC_NT_ switch, the circular variances *V*(*ϕ*) and *V*(*ψ*) [47] of the backbone dihedral angles *ϕ* and *ψ* have been calculated (Eq. 1 in Materials and Methods). In the center of the belt, the *α* helices 485-496 (A1) and 465-471 (E1) observed in the X-ray crystallographic structures (Figure 1A) display minimum values of *V*(*ϕ*) and *V*(*ψ*) for most of the trajectories, whereas peaks of fluctuations are located mostly in the flanking outside regions (Figure 9). The open state of A1 shows the largest interval of null values, corresponding to the most stable *α* helix. Overall, the closed states display shorter ranges of minimal circular variances in *α* helix. For closed states (orange, green, brown and olive green curves), as well as for E1ope70 and E1ope47 (magenta and cyan curves), the acidification induces appearance of peaks inside and outside of the *α* helix. This increased belt mobility could be a starting point for belt destabilization at acidic pH before translocation.

**Figure 9.**
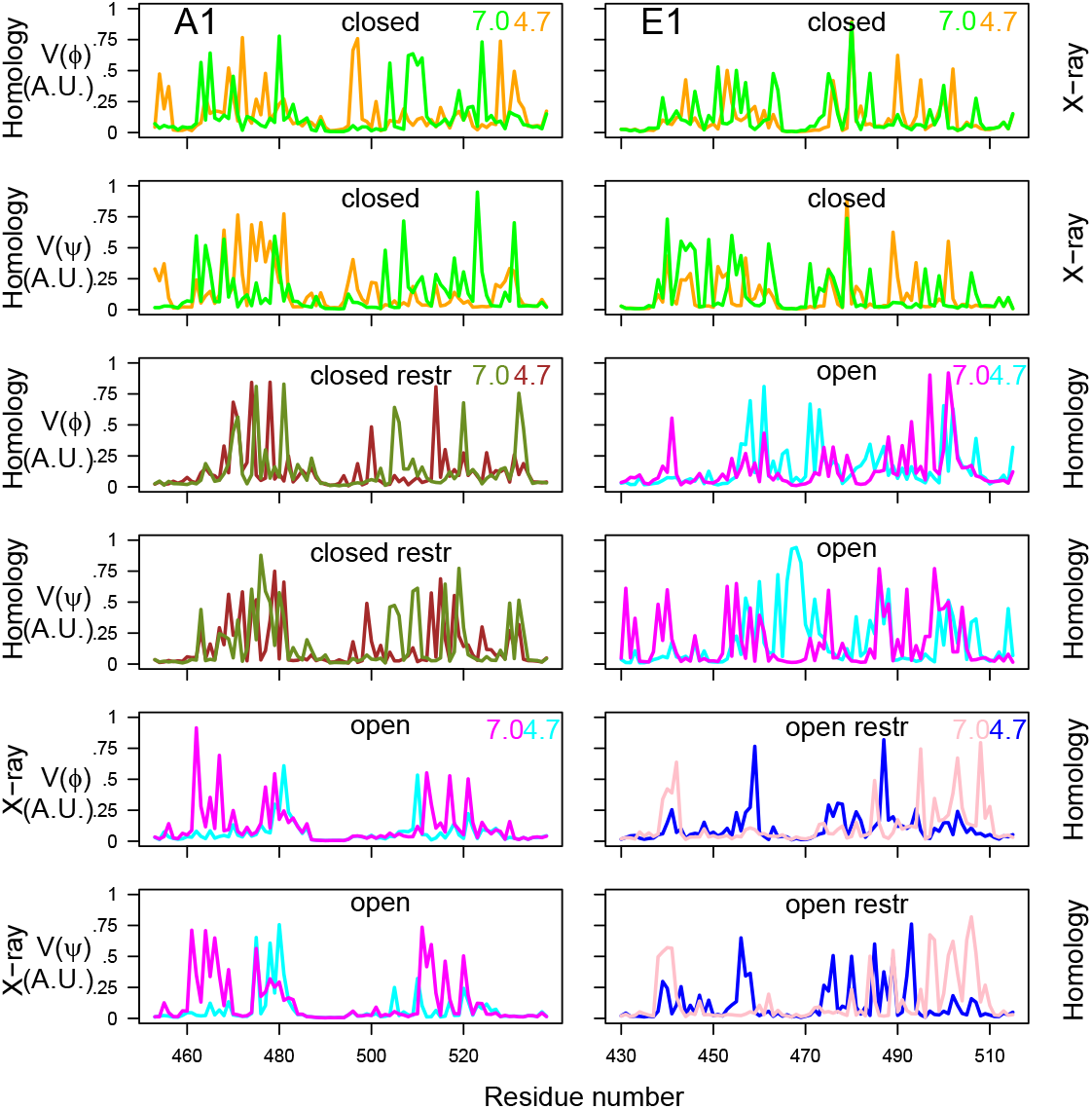
Variations of circular variances *V*(*ϕ*) and *V*(*ψ*) (Eq. 1) [47] calculated on the interval 150-300 ns of the trajectories, for dihedral angles *ϕ* and *ψ* of belt domain. The color code is the following: cyan (A1ope47, E1ope47), blue (E1ope47r), magenta (A1ope70,E1ope70), pink (E1ope70r), orange (A1clo47, E1clo47), brown (A1clo47r), green (A1clo70, E1clo70), olive green (A1clo70r). The pH values used to define the protonation level of residues are written on the right part of the plots. The title of each plot refers to the BoNT type (A1/E1), to the conformational state (open/closed). The “open restr” and “closed restr” titles correspond to the trajectories A1clo47r, A1clo70r, E1ope47r and E1ope70r, in which restraints have been used during the homology modeling step (Table S3).

The circular variance calculated on HC_NT_ switch (Figure 10) most often displays a peak of mobility in the middle of this region, spanning residues 635-640 and 648-652 for BoNT/A1 and residues 610-618 for BoNT/E1. For the two BoNTs, the region of maximum variance includes the loop between the helices *α_A_* and *α_B_* of the HC_NT_ switch, these helices names having been proposed by Lam et al [30]. This loop is also the region displaying the largest conformational transition in [30].

**Figure 10.**
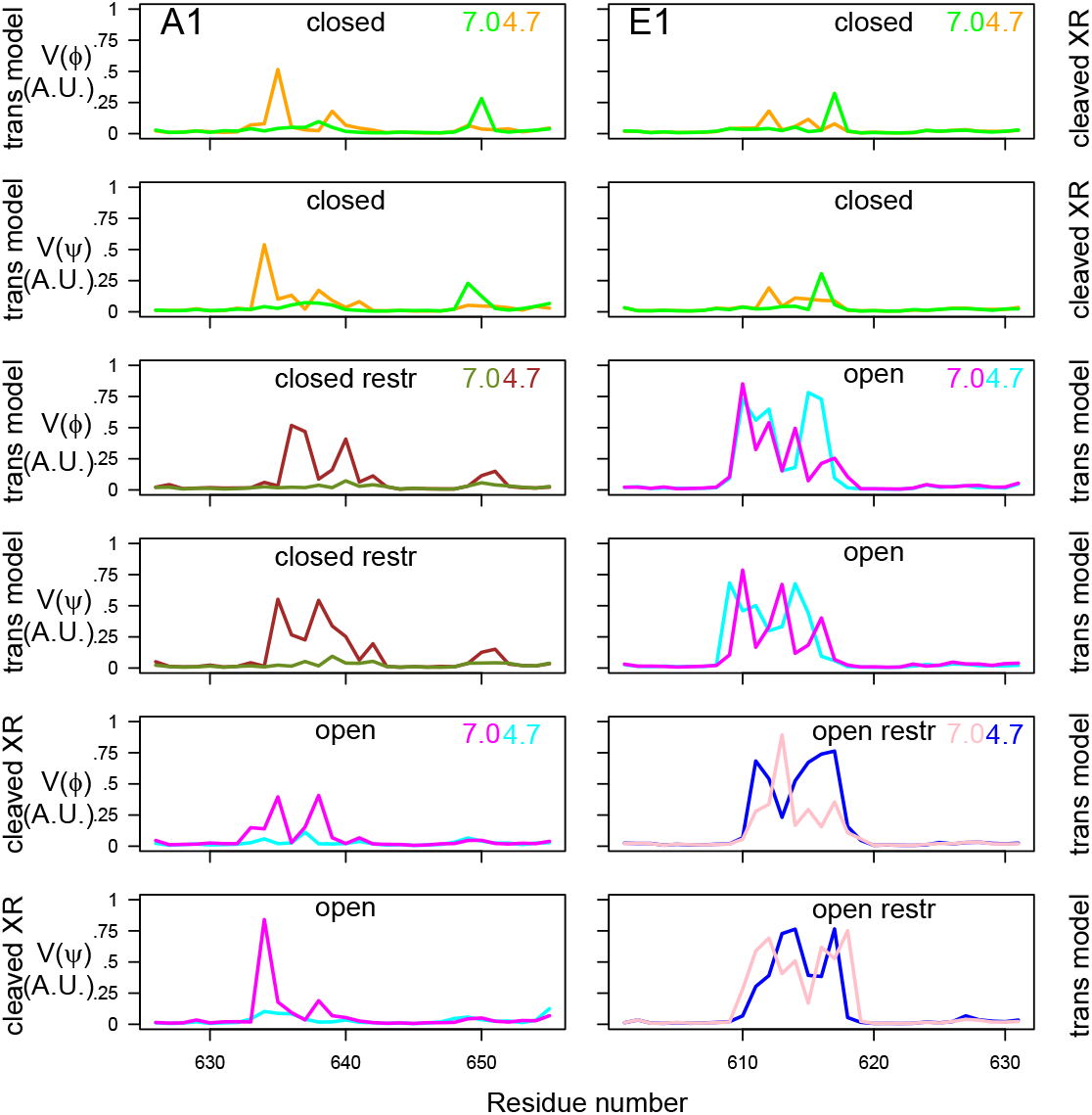
Variations of circular variances *V*(*ϕ*) and *V*(*ψ*) (Eq. 1) [47] calculated on the interval 150-300 ns of the trajectories, for dihedral angles *ϕ* and *ψ* of the switch domain in HC_NT_. The color code as well as the titles and annotations are the same than in Figure 9.

The circular variance in HC_NT_ switch display different features for the various trajectory conditions: in the closed state of BoNT/A1, the mobility increases at acidic pH, whereas it decreases at acidic pH for the open state of BoNT/A1. In BoNT/E1, no pH effect is observed.

#### 2.4.3. HC_NT_ helices bending

The bending of *α* helices located in HC_NT_ was analyzed using Bendix [48] (Figure 11). In the *α* helices 1 and 2, the maxima of bending angles are located in the residue ranges 705-715 and 777-797 (A1) and 698-718 and 737-798 (E1). This corresponds to a bend already observed in the X-ray structures and initial models, located slightly at the bottom of the HC_NT_ switch. The local bending angles of *α* helices 1 spanning residues 680-740 (A1) and 660-720 (E1) display similar profiles for all conditions with peaks of bending at the middle of the helix (Figure 11). On the other hand, the helices 2 spanning residues 760-820 (A1) and 740-800 (E1) display variations of bending angles depending on the closed or open state and on the type of BoNT. In BoNT/E1, the bending peaks are located around residue K^775^, located close to several protonated residues (section “Analysis of the protonation at varying pH in different states”).

**Figure 11.**
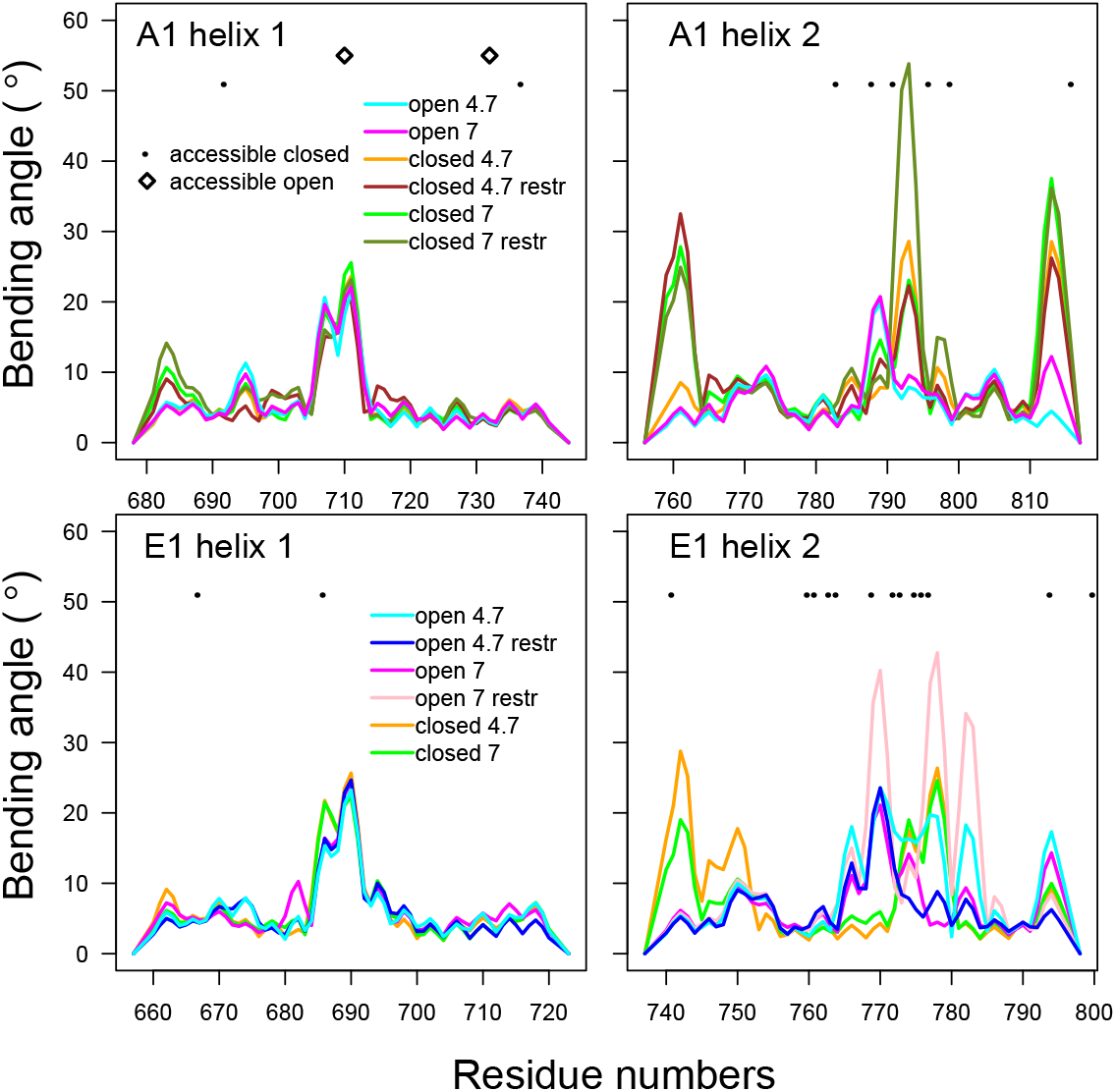
Analysis of the bending angles of the helices in the domain of translocation of BoNTs A1 and E1. Local bending angles were calculated using the Bendix VMD plugin [48] on the *α* helices 678-744 and 756-817 in A1, and on the *α* helices 657-723 and 737-798 in E1. The profiles of these angles are drawn with the color curves. The color code of the curves is the following: cyan (A1ope47, E1ope47), blue (E1ope47r), magenta (A1ope70,E1ope70), pink (E1ope70r), orange (A1clo47, E1clo47), brown (A1clo47r), green (A1clo70, E1clo70), olive green (A1clo70r). In the same plots residues that show a difference in solvent accessible surface are indicated by different symbols depending on whether the accessible surface is larger in the trajectory starting from closed (•) or open (◇) conformations.

To summarize, complicated correlation patterns are observed between the destabilization of the *α* helix belt, the more accessible residue surfaces in HC_NT_, the residue protonation, the bending of HC_NT_ helices and the internal mobility of the HC_NT_ switch. In several cases at acidic pH and in closed states, more accessible surfaces as well as flexibility in belt and HC_NT_ switch are observed simultaneously. The observation of larger accessible surfaces at acidic pH agrees with the recent molecular dynamics study of BoNT/E1 [35]. The results of the present work will be discussed in the next section.

## 3. Discussion

The relevance of BoNTs proteins in the medical field has stimulated in recent years the formulation of mechanistic hypotheses accounting for variations in their kinetics and stability. However, the sequence of events at the basis of their function is far from being understood at the molecular level, although numerous structures of the interaction partners have been determined. Some key unanswered questions about BoNTs are: first, how ternary and quaternary structural changes of the protein relate to the relevant physiological processes? second, could we capture structural signatures underlying pH-induced mechanisms? third, could molecular level based knowledge of the action of different BoNTs guide structure-based engineering for toxin-based therapeutics?

Addressing these points requires at first reliable, atomically refined structures of the protein complexes, and an extensive knowledge of the network of residue-residue interactions, which unfortunately is currently available as X-ray or CryoEM maps only for selected BoNT subtypes in particular conformations engineered as single chains. In any event, the protein structure-dynamics-function paradigm requires the knowledge of how do proteins dynamically behave in water and/or in the presence of a membrane.

The aim of this work is to provide insight on the different possible conformations of BoNTs and on how these are affected by the environment, via computational tools. Two BoNT subtypes, A1 and E1, have been studied via full atomistic molecular dynamics simulations corresponding to a cumulative trajectory duration of 3.6 *μ*s, both in the open and closed conformations. Two different protonation states have been considered corresponding to acidic and neutral pH values of respectively 4.7 and 7.0.

Different initial structures have been exploited, starting from cleaved X-ray crystallographic conformations as well as from trans models, based on sequence alignment between BoNT/A1 and BoNT/E1. Given that the two studied toxins, A1 and E1, display sequences identity of about 35 and 45%, depending on the chain, we cannot affirm that trans models correspond to conformations indeed significantly populated in the conformational landscape.

Results provide a structural and functional annotation of full-length BoNTs composed by two distinct protein chains, which agrees to the molecular dynamics study of BoNT/E1 at various pH values [35]. A global overview of the simulations results shows that movement apart of domains prevails on the tertiary rearrangement, in agreement with the results of Chen et al [33] on BoNT/A. The parallel use of different starting points permits the detection of conformational features which can be related to the BoNT function, as movement apart of domains, internal flexibility of HC_CC_ ganglioside binding site, of HC_NT_ switch and of the belt, and the larger accessibility to solvent of residues in the HC_NT_ and LC domains. Moreover the data pointed out to connections between different regions, as the belt and the HC_NT_ domains, or the HC_NT_ domain and the HC_CN_ and HC_CC_ domains. Given that most of these observations can be related to independent experimental observations, they provide insights on the functional dynamics of BoNTs. In particular, residues displaying larger accessible surfaces in the translocation domain HC_NT_ could be starting anchors for the interaction of BoNTs with the membrane.

Remarkably, the BoNT/E1, simulated at pH 7, displayed a large drift from the starting X-ray crystallographic structure. This supports a picture of the BoNT/E1 structure in solution in which the domains LC and HC_CN_/HC_CC_ spontaneously move apart from each other. One should also notice that HC/E takes various positions with respect to LC in the structures of the *C. botulinum* progenitor M complex of type E [49,50]; this agrees with an internal mobility of BoNT/E1 which is able to occupy conformations different from the closed one.

The variations of residue protonation due to pH have some effects, although of minor extent compared with the results from experiments conducted in the presence of membrane [20,21,51]. At acidic pH and in closed state, a patch of HC_NT_ residues close in 3D space is more exposed to the solvent. The observation of this patch correlates with the largest number of non-histidine residues protonated in HC_NT_. The internal mobility of belt, and in particular of belt *α* helix, allows one to propose a model for the initiation of translocation, in which the larger mobility of belt is transmitted to HC_NT_ through the connection loop, inducing a more favorable interaction of HC_NT_ residues with the non-polar membrane environment.

A recent article has described structures of BoNT/B and BoNT/E obtained by Cryo-electron microscopy (Cryo-EM) [31]. Several observations reported are in good agreement with the data presented here. First, the main structural variations seem to come from overall movement of the domains with respect to each other, similarly to the observations made here that the RMSD of individual domains (Figure 3) are smaller than the global RMSD values (Figure 2). The binding domain in BoNT/B shifts up to 2 Å around HC_CN_ and this seems to be further accentuated at the HC_CC_ domain [31]. This observation agrees with the increase of distance between the domains HC_NT_ and HC_CN_/HC_CC_ here observed (Figure 5). In addition, the EM map quality around the binding domains HC_CN_ and HC_CC_ is generally weaker compared to the rest of the toxin and the map is particularly well-defined for LC, in agreement with the RMSD and RMSF profiles observed in MD trajectories (Figures 2 and 4) and with the mobility observed here for the lipid-binding loop (LBL). In the cryo-EM map of BoNT/B, the belt is well-ordered except for a small surface-exposed *α*-helix, which is the one for which we observed variations in internal mobility (Figure 9). More generally, results from the present work confirm the fact that the X-ray crystallographic structures of BoNTs, determined from samples formed from one chain, do not capture the structural features of the toxin in solution, which plays an essential role into the physico-chemical aspects underlying the functional physiological processes. This observation agrees with recent work [44] showing that BoNT/B in interaction with the receptors seems to display much more molecular flexibility that what can be deduced from the X-ray crystallographic structures of BoNTs.

## 4. Materials and Methods

### 4.1. Preparation of starting conformations of toxins

In order to prepare systems in which BoNTs are formed by two protein only connected by a disulfide bridge, the structures from PDB entries 3BTA (A1) [7] and 3FFZ (E1) [9] have been cleaved, eliminating the peptides ^438^TKSLDKGYNK^447^ and ^420^GIR^422^, the numbering being taken from PDB files 3BTA and 3FFZ (MR Popoff, personal communication). Consequently, the BoNT domains are defined in the present work from ranges of residues given in Table 2. After removing the cleaved sequences, the disulfide bridge is established between C^430^ and C^444^ in BoNT/A1 (Figure 1D) and between C^412^ and C^423^ in BoNT/E1.

For the two studied BoNTs, the conformations of the open state were prepared from the cleaved 3BTA structure and the conformations of the closed state using the cleaved 3FFZ structure. These starting points of the simulations were refined using Modeller [52,53] to provide an energy optimization of the structures and to relax any structural stress due to the cleavage in two chains. These two conformations, corresponding to the open state of BoNT/A1 and to the closed state of BoNT/E1, will be denoted in the present work as cleaved X-ray models.

The closed A1 and open E1 conformations of BoNTs were built by homology modeling using Modeller [52,53] and the sequence alignments obtained by T-Coffee [54] (Figure S7 and S8) The percentages of identity between the sequences of the two toxins are of 35% for LC and of 43% for HC, for which the use of the homology approach is valid [55]. The template structures used for homology modelling were the PDB entries 3BTA for the open state, and 3FFZ for the closed state, cleaved as described above. An additional homology model was built for each BoNT by imposing restraints (Table S3) between the *α* helix 485-496 (A1) or 465-471 (E1) in the belt region (Figure 1A) and the other domains of BoNTs, in order to stabilize the helix and hopefully the belt conformation. These four starting points for the simulations obtained by homology (two without and two with restraints, respectively) will be denoted trans models.

For each Modeller run, 200 conformations were generated using the modeling refinement method “very_slow” [52]. The conformation displaying the best score DOPE was selected, the hydrogens were added to the models and the flip of sidechains optimized by Molprobity [56]. The final models were then evaluated using Molprobity and QMEAN [57] scores (Table S4) The quality scores of the models are similar to the ones obtained for the initial X-ray crystallographic structures 3BTA and 3FFZ. Surprisingly the scores are slightly better for trans models than for X-ray cleaved models.

Starting from the previously calculated models, the protonation of residues at neutral (7.0) and acidic (4.7) was predicted using the web-based implementation of H++, a method for pKa calculations based on continuum electrostatic model [58,59]. The list of protonated residues are given in Tables S1 and S2, along with the value of the midpoint pK(1/2) of the titration curve predicted. In the majority of cases the latter can be approximated by the classical sigmoidal (Henderson-Hasselbalch) shape, in which case pK(1/2) = pKa [60]. The protonation state of each titrable residue is then assigned by H++ basing on the comparison between pK(1/2) and the selected pH value. In detail, if pK(1/2) is ≥ pH the residue is considered as protonated. A detailed discussion on the accuracy and benchmarks of H++ predictions is contained in refs. [58,59].

Using the two different protonation levels on the six previously defined systems, twelve systems were finally set-up. They were named (Table 1) using the type of BoNT (A1/E1), the conformational state (clo for close, ope for open) and the pH for which the protonation was defined (47 or 70). The suffix “r” was added in the case where restraints from Table S3 were used during the homology modelling. The names of these systems will be also the names of the corresponding trajectories.

### 4.2. Molecular dynamics simulations

For each previously described system, the protein was embedded in a large water box (183×148×194 Å), and chloride counterions were added to neutralize the net system charge. The total number of atoms was about 510000 in each case, see Table 1 for the details on the systems’ composition. All molecular dynamics (MD) simulations were performed using NAMD 2.13 [61], with the CHARMM(Chemistry at Harvard Macromolecular Mechanics)36 force field [62] for protein and the TIP3P (Transferable Intermolecular Potential with 3 Points) model for water [63]. A cutoff of 12 Å and a switching distance of 10 Å were used for non-bonded interactions, while long-range electrostatic interactions were calculated with the Particle Mesh Ewald (PME) method [64]. The RATTLE algorithm [65] was used to keep rigid all covalent bonds involving hydrogen atoms, enabling a time step of 2 fs. At the beginning of each trajectory, the system was minimized for 20000 steps, then heated up gradually from 0 K to 310 K in 31000 integration steps. Finally, the system was equilibrated for 50.000 steps in the NVT ensemble at 310K. In these first three stages, all carbon *α* atoms were kept fixed. Simulations were then performed in the NPT ensemble, (P= 1bar, T=310K), with all atoms free to move. Atomic coordinates were saved every 10 ps. Protein roto-translation motions were harmonically restrained throughout the simulations (with a scaled force constant of 10 kcal/mol), to avoid any rigid-body protein motion but leaving unaltered the internal dynamics. For each trajectory, 300 ns of production were recorded for a cumulative trajectory duration of 3.6 *μ*s.

### 4.3. Analysis of molecular dynamics trajectories

The analysis of BoNT conformations sampled along MD trajectories was realized using ccptraj [66] and the python package MDAnalysis [37,38]. The solvent accessible surfaces were calculated using FreeSASA on every 10 frames of the trajectory[67]. The solvent accessible surfaces of residues were averaged along the time interval 150-300 ns in each of the six trajectories of Table 1, producing for each residue, six values: *S*_*clo*47_, *S*_*clo*47*r*_, *S*_*clo*70_, *S*_*clo*70*r*_, *S*_*ope*47_, *S*_*ope*70_ for BoNT/A1, and *S*_*clo*47_, *S*_*clo*70_, *S*_*ope*47_, *S*_*ope*47*r*_, *S*_*ope*70_, *S*_*ope*70*r*_ for BoNT/E1. For each BoNT and each residue, six normalized surface values were obtained by dividing them by the residue surface *S_ave_* equal to 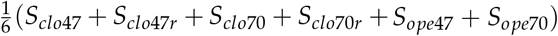 for BoNT/A1 and equal to 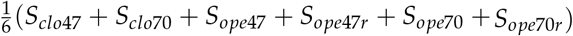 for BoNT/E1. The obtained ratio values smaller than 0.9 (respectively larger than 1.1) were pointed out to correspond to smaller (respectively larger) accessible surfaces with respect to *S_ave_*.

The bending angles of *α* helices 1 and 2 (Figure 1B) in HC_NT_ were determined using the VMD plugin Bendix [48] on every 10 frames of the trajectory. The figures containing structures were prepared using VMD [68] or PyMOL [69].

The relative variability of backbone angles *ϕ* and *ψ* in the belt domain was estimated by calculating the circular variances *V*(*ϕ*) and *V*(*ψ*). For a given angle *θ*, this parameter is defined as:

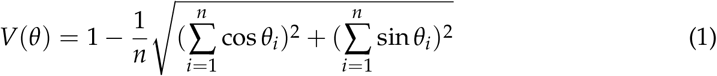

where *n* is the number of trajectory frames considered and *θ_i_* is the angle value in the frame *i*.

## Supplementary Materials

The following supporting information can be downloaded at: www.mdpi.com/xxx/s1, Figure S1: Comparison of the first frame and of the frame recorded at 126.6ns for the trajectory E1clo70; Figure S2: Structure of the A1 HC_CC_ domain in the X-ray structure 3BTA, and along the MD simulation A1ope70 (initial and last frame). Figure S3: Position along the protein sequence of the four sets of residues displaying similar variations of the solvent accessible surface along the MD; Figures S4-S6: Subsets of residues with similar variations of the solvent accessible surfaces displayed on the protein structure. Figure S7: Alignment of the BoNT/A1 and BoNT/E1 sequences from the LC chain; Figure S8: Alignment of the BoNT/A1 and BoNT/E1 sequences from the HC chain; Table S1: List of protonated residues in BoNT/A1; Table S2: List of protonated residues in BoNT/E1. Table S3: Restraints applied during the Modeller run; Table S4: Quality parameters for the starting points of simulation. Zip archive representative_conformations.zip: contains representative conformations extracted from the molecular dynamics trajectories using the **S**elf-**O**rganizing **M**aps (SOM), as described in the Supplementary Materials.

## Author Contributions

TEM, GC and LC prepared the systems, performed the molecular dynamics simulations and analyzed the trajectories. TEM and GC wrote the manuscript with the help of CRE. EL and MRP put the introduction into microbiology perspective. GC, LC, LM, MRP, CRE, EL and TEM reviewed the manuscript.

## Data Availability Statement

The data presented in this study are openly available at 10.5281/zen-odo.6989010.

