## Supplementary Material for "*In silico* conformational features of botulinum toxins A1 and E1 according to the intraluminal acidification"

### Extracting representative conformations from molecular dynamics trajectories

Representative conformations were extracted from each of the molecular dynamics (MD) trajectories listed in Table 2 of the main text, using a clustering approach, the **S**elf-**O**rganizing **M**aps (SOM). SOM is an artificial neural network (ANN) trained using unsupervised learning [1]. The conformations sampled along trajectories were encoded from the distances $d_{ij}$ calculated between the $n$ $C_\alpha$ atoms of BoNT, by diagonalizing the covariance matrix $C$:

$$C_{i,j} = \frac{1}{n}\sum_{k=1}^{n}\sum_{l=1}^{n}(d_{i,k} - \bar{d}_i)(d_{l,j} - \bar{d}_j) \quad (1)$$

where $\bar{d}_i = \frac{1}{n}\sum_{j=1}^{n} d_{i,j}$. The information contained in the matrix $C$ is equivalent to its four largest eigenvalues along with the corresponding eigenvectors. The eigenvalue and eigenvector descriptors are used to train a periodic Euclidean 2D self-organizing map (SOM), defined by a three-dimensional matrix. The self-organizing maps were initialized with a random uniform distribution covering the range of values of the input vectors. At each step, an input vector is presented to the map, and the neuron closest to this input is updated. The maps are trained in two phases. During the first phase, the input vectors are presented to the SOM in random order to avoid mapping bias with a learning parameter of 0.5, and a radius parameter of 36. During the second phase, the learning and radius constants are decreased exponentially from starting values 0.5 and 36, respectively, during 10 cycles of presentation of all the data in random order. Once the calculation of the SOM has been realized, the processed MD trajectory has been transformed into a map in which each pixel corresponds to a series of similar conformations of BoNT. The conformations of BoNT corresponding to local maxima of homogeneity in the SOM map, are representative conformations of the MD trajectory and are made available in the archive representative_conformations.zip

**Figure S1.** Comparison of the first frame and of the frame recorded at 126.6ns for the trajectory E1clo70. The BoNT/E1 conformations are drawn in cartoon, and colored in the following way: LC (green), belt (cyan), $HC_{NT}$ (orange), $HC_{CN}$ (magenta) and $HC_{CC}$ (red).

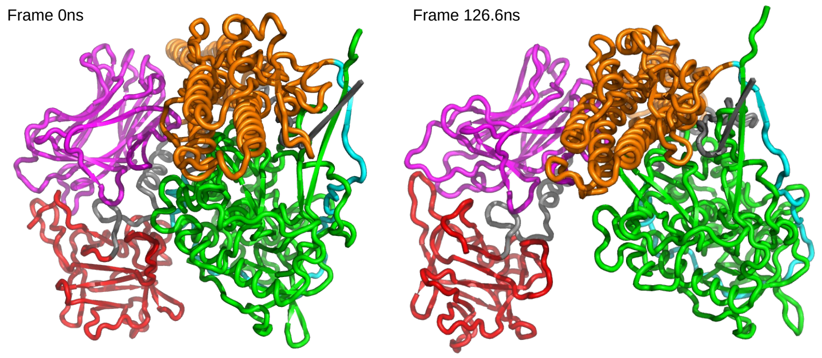

**Figure S2.** Structure of the $HC_{CC}$ domain in the A1 open X-ray structure (PDB entry: 3BTA), in the initial frame and the final (300ns) frame of the A1ope70 trajectory. The protein is drawn in cartoon; the LBL (residues 1188-1198), the GBS (residues 1255-1257) and the residue stretch 1252-1254 are also represented as surface in yellow, orange and magenta, respectively, to convey the information on the steric hindrance of aminoacids.

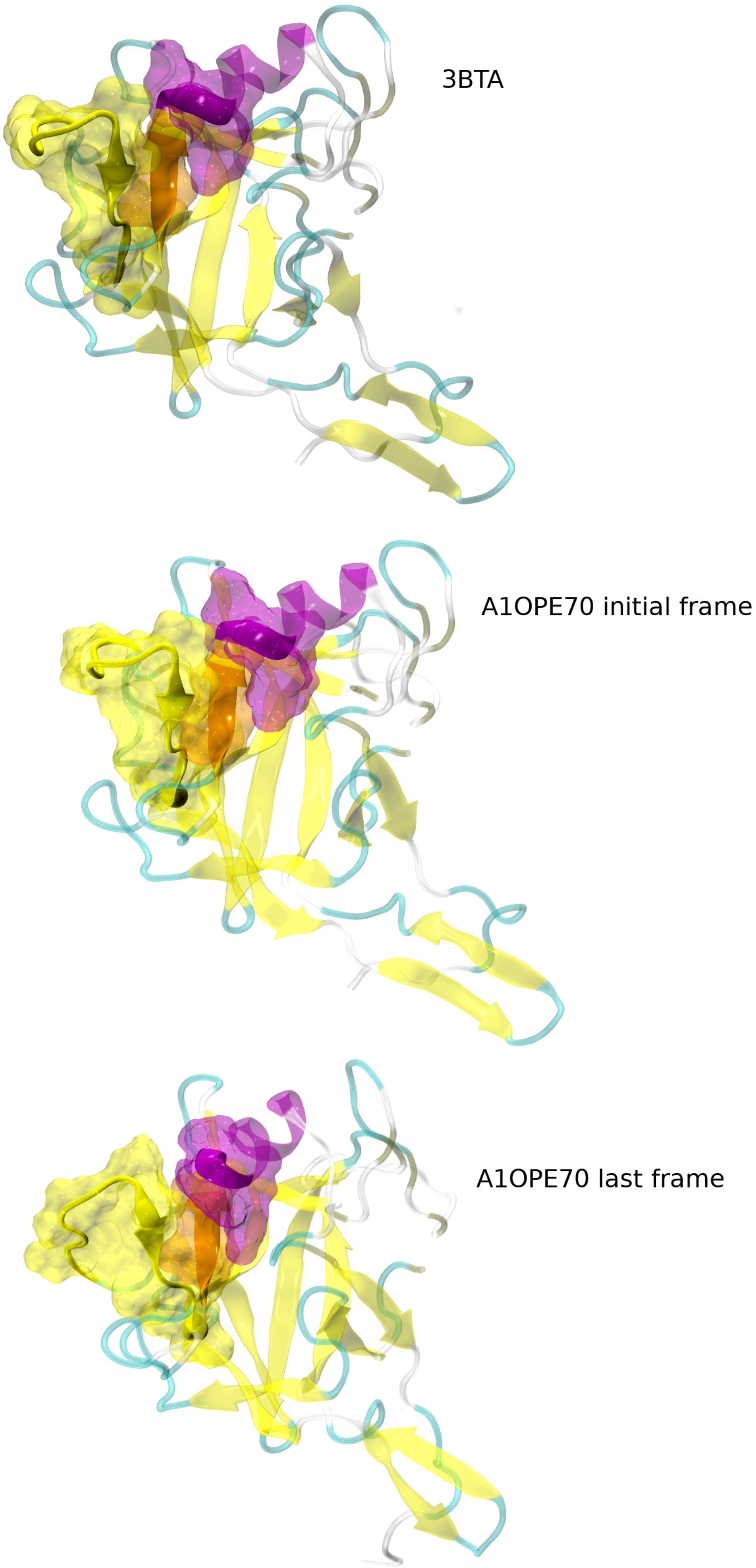

**Figure S3.** Sets of residues displaying similar variations of surfaces accessible to solvent along the MD trajectories recorded on the botulinum toxins A1 and E1. The dashed lines mark domain boundaries.

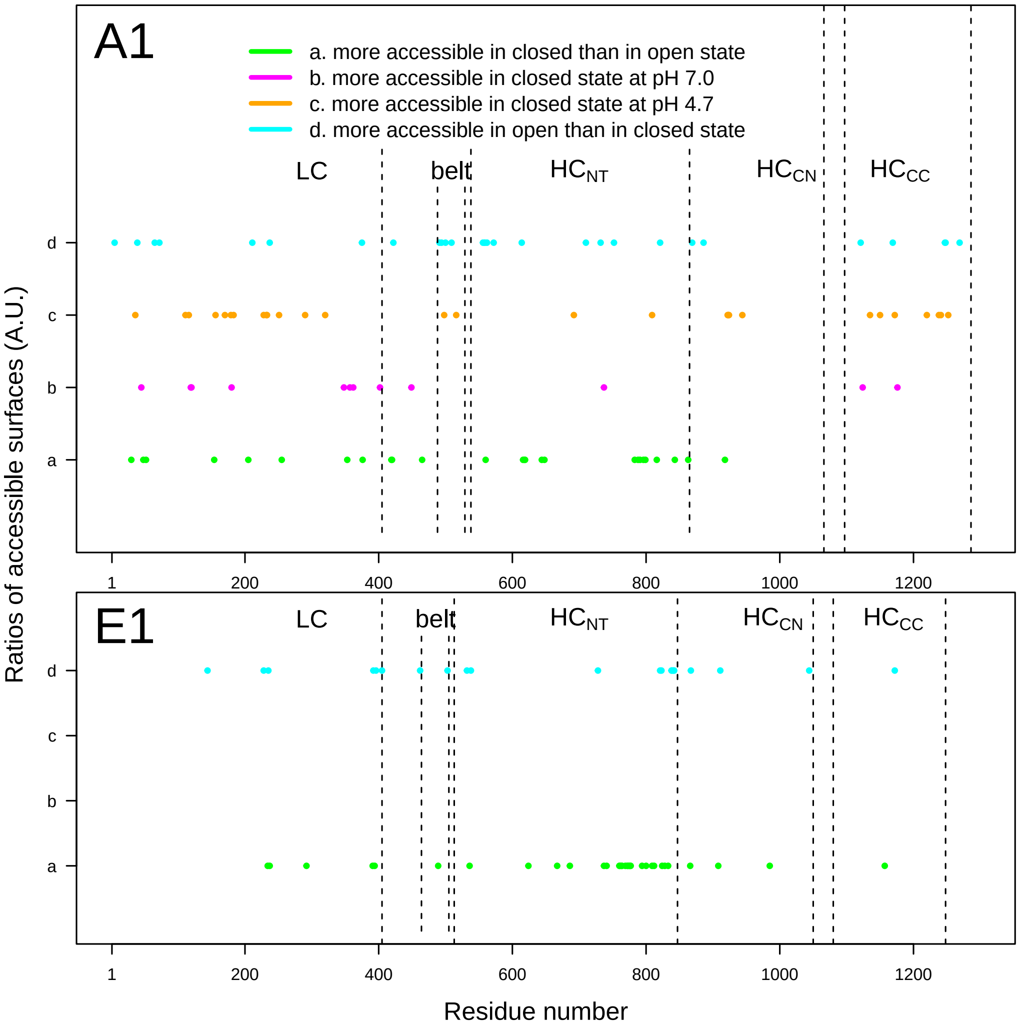

**Figure S4.** View of the $HC_{NT}$ domain in cartoon for BoNT/A1 (top) and BoNT/E1 (bottom). The switch and the C terminal $\alpha$ helix are colored in orange, and the $\alpha$ helix 2 is colored in yellow. The residues more accessible in closed than in open state are drawn in licorice and colored in green.

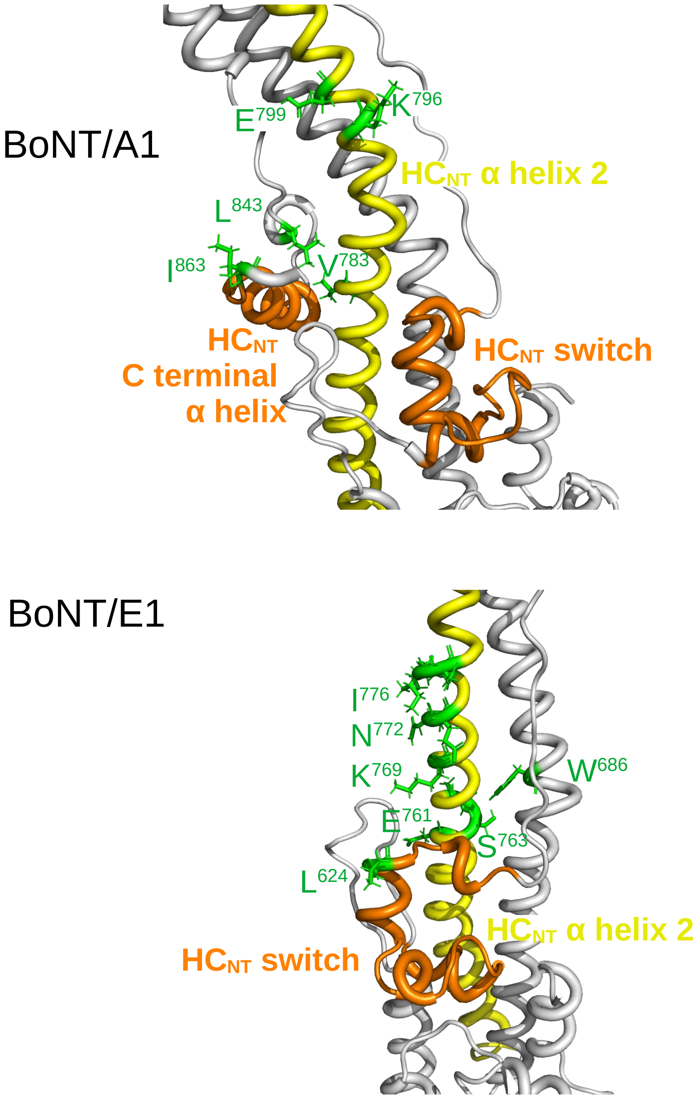

**Figure S5.** View of the LC domain in cartoon for BoNT/A1 (top) and BoNT/E1 (bottom). The residues more accessible in closed than in open state are drawn as van der Waals spheres and colored in green, whereas the residues protonated at acidic pH are drawn in licorice and colored in orange.

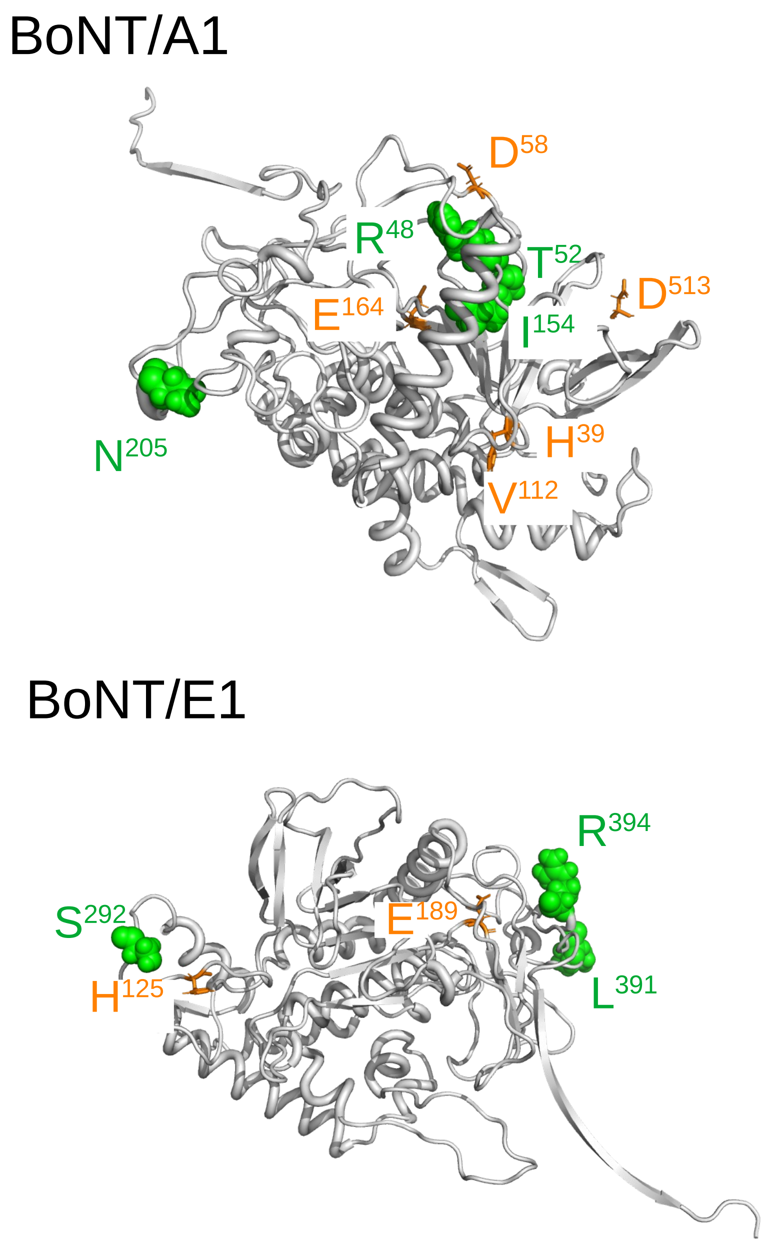

**Figure S6.** View of the LC domain in cartoon for BoNT/A1. The residues more accessible in closed than in the open state at acidic pH are drawn in licorice and colored in green. The residues H223 and H227 of the catalytic site are drawn in licorice and colored in orange.

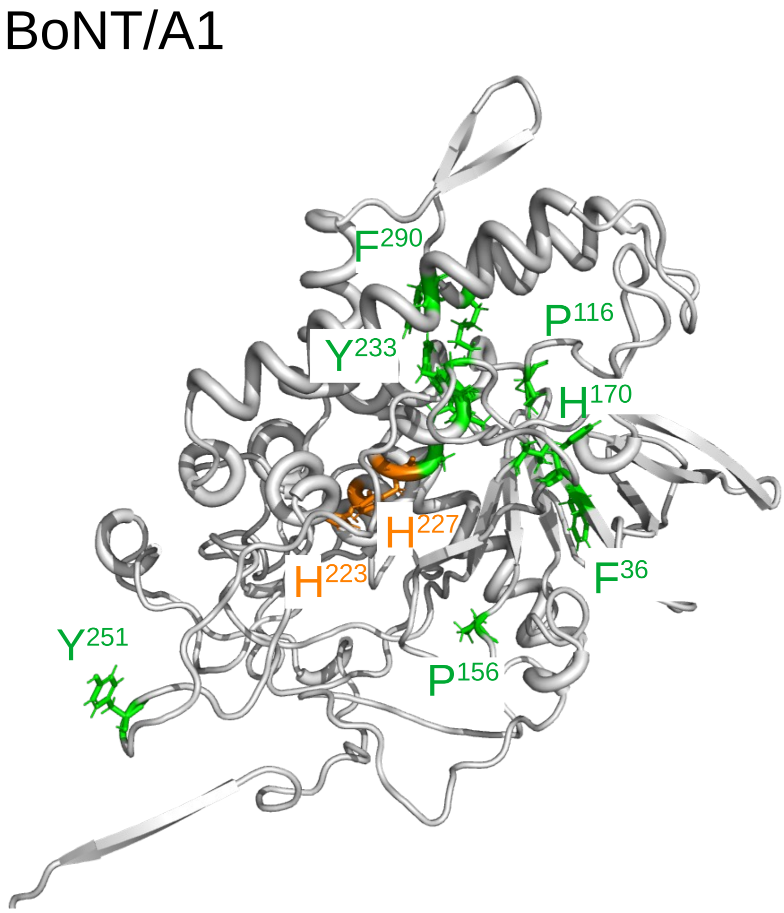

**Figure S7.** Alignment of the BoNT/A1 and BoNT/E1 sequences from the LC chain obtained with T-Coffee version 9.01 [2]. The percentage of sequence identity is 35%.

```
A1_3BTA_LC      MPFVNKQFNYKDPVNGVDIAYIKIPNAGQMQPVKAFKIHNKIWVIPERDT
E1_LC           MPKIN-SFNYNDPVNDRTILYIKP--GGCQEFYKSFNIMKNIWIIPERNV
                ** :* .***:****.  * ***   .*  :  *:*:* ::**:****:.
A1_3BTA_LC      FTNPEEGDLNPPPEAKQVPVSYYDSTYLSTDNEKDNYLKGVTKLFERIYS
E1_LC           IGTT-PQDFHPPTSLKNGDSSYYDPNYLQSDEEKDRFLKIVTKIFNRINN
                : ..   *::**.. *:   ****..**.:*:***.:** ***:*:** .
A1_3BTA_LC      TDLGRMLLTSIVRGIPFWGG-STIDTELKVIDTNCINVIQPDGSYRSEEL
E1_LC           NLSGGILLEELSKANPYLGNDNTPDNQFHIGDASAVEIKFSNGSQDILLP
                .  * :** .: :. *: *. .* *.:::: *:..:::  .:**
A1_3BTA_LC      NLVIIGPSADIIQFECKSFGHEVL-NLTRNGYGSTQYIRFSPDFTFGFEE
E1_LC           NVIIMGAEPDLFETNSSNISLRNNYMPSNHRFGSIAIVTFSPEYSFRFND
                *::*:*...*::: :...:. .     :.: :**   : ***:::* *::
A1_3BTA_LC      SLEVDTNPLLGAGKFATDPAVTLAHELIHAGHRLYGIA-INPNRVFKVNT
E1_LC           NC---------MNEFIQDPALTLMHELIHSLHGLYGAKGITTKYTITQKQ
                .           .:*  ***:** *****: * ***   *..: .:. :
A1_3BTA_LC      NAYYEMSG-LEVSFEELRTFGGHDAKFIDSLQENEFRLYYYNKFKDIAST
E1_LC           NP--LITNIRGTNIEEFLTFGGTDLNIITSAQSNDIYTNLLADYKKIASK
                *.   ::.   ..:**: **** * ::* * *.*::      .:*.***.
A1_3BTA_LC      LNKAKSIVGTTASLQYMKNVFKEKYLLSEDTSGKFSVDKLKFDKLYKMLT
E1_LC           LSKVQV---SNPLLNPYKDVFEAKYGLDKDASGIYSVNINKFNDIFKKLY
                *.*.:    :.. *:  *:**: ** *.:*:** :**:  **:.::* *
A1_3BTA_LC      EIYTEDNFVKFFKVLNRKTYLNFDKAVFKINIVPKVNYTIYDGFNLRNTN
E1_LC           S-FTEFDLRTKFQVKCRQTYIGQYKYFKLSNLLNDSIYNISEGYNIN--N
                . :** :: . *:*  *:**:.  * .   *:: .  *.* :*:*:.  *
A1_3BTA_LC      LAANFNGQNTEINNMNFTKLKNFTGLFEFYKLLCVRGIITSKTKSLDKGY
E1_LC           LKVNFRGQNANLNPRIITPITG-------------RGLVKKIIRFCKNIV
                * .**.***:::*   :* :..             **::..  :  .:
A1_3BTA_LC      NKA
E1_LC           SVK
                .
```

**Figure S8.** Alignment of the BoNT/A1 and BoNT/E1 sequences from the HC chain obtained with T-Coffee version 9.01 [2]. The percentage of sequence identity is 43%.

```
A1_3BTA_HC      -------NNWDLFFSPSEDNFTND-LNKGEEITSDTNIEAAEENISLDLI
E1_HC           KSICIEINNGELFFVASENSYNDDNINTPKEIDDTV---TSNNNYENDL-
                       ** :*** .**:.:.:* :*. :** . .   ::::* . **
A1_3BTA_HC      QQYYLTFNFDNEPENISIENLSSDIIGQLELMPNIER--FPNGKKYELDK
E1_HC           DQVILNFNSESAP-GLSDEKLNLTIQND-AYIPKYDSNGTSDIEQHDVNE
                :*  *.** :. * .:* *:*.  * .:   :*: :    .: :::::::
A1_3BTA_HC      YTMFHYLRAQEFEHGKSRIALTNSVNEALLNPSRVYTFFSSDYVKKVNKA
E1_HC           LNVFFYLDAQKVPEGENNVNLTSSIDTALLEQPKIYTFFSSEFINNVNKP
                 .:*.** **:. .*:..: **.*:: ***: .::******:::::***.
A1_3BTA_HC      TEAAMFLGWVEQLVYDFTDETSEVSTTDKIADITIIIPYIGPALNIGNML
E1_HC           VQAALFVSWIQQVLVDFTTEANQKSTVDKIADISIVVPYIGLALNIGNEA
                .:**:*:.*::*:: *** *:.: **.******:*::**** ******
A1_3BTA_HC      YKDDFVGALIFSGAVILLEFIPEIAIPVLGTFALVSYIA---NKVLTVQT
E1_HC           QKGNFKDALELLGAGILLEFEPELLIPTILVFTIKSFLGSSDNKNKVIKA
                 *.:* .** : ** ***** **: **.: .*:: *::.   **  .:::
A1_3BTA_HC      IDNALSKRNEKWDEVYKYIVTNWLAKVNTQIDLIRKKMKEALENQAEATK
E1_HC           INNALKERDEKWKEVYSFIVSNWMTKINTQFNKRKEQMYQALQNQVNAIK
                *:***.:*:***.***.:**:**::*:***::  :::* :**:**.:* *
A1_3BTA_HC      AIINYQYNQYTEEEKNNI--NFNIDDLSSKLNESINKAMININKFLNQCS
E1_HC           TIIESKYNSYTLEEKNELTNKYDIKQIENELNQKVSIAMNNIDRFLTESS
                :**: :**.** ****::  :::*.::..:**:.:. ** **::**.:.*
A1_3BTA_HC      VSYLMNSMIPYGVKRLEDFDASLKDALLKYIYDNRGTLIGQV-DRLKDKV
E1_HC           ISYLMKIINEVKINKLREYDENVKTYLLNYIIQ-HGSILGESQQELNSMV
                :****: :    :::*.::* .:*  **:** : :*:::*:  :.*:. *
A1_3BTA_HC      NNTLSTDIPFQLSKYVDNQRLLSTFTEYIKNIINTSILNLRYESNHLIDL
E1_HC           TDTLNNSIPFKLSSYTDDKILISYFNKFFKRIKSSSVLNMRYKNDKYVDT
                .:**...***:**.*.*:: *:* *.:::*.* .:*:**:**:.:: :*
A1_3BTA_HC      SRYASKINIGSKVNFDPIDKNQIQLFNLESSKIEVILKNAIVYNSMYENF
E1_HC           SGYDSNININGDVYKYPTNKNQFGIYNDKLSEVNISQNDYIIYDNKYKNF
                * * *:***...*   * :***: ::* : *::::  :: *:*:. *:**
A1_3BTA_HC      STSFWIRIPKYFNSI-SLNNEYTIINCM-ENNSGWKVSLNYGEIIWTLQD
E1_HC           SISFWVRIPNYDNKIVNVNNEYTIINCMRDNNSGWKVSLNHNEIIWTFED
                * ***:***:* *.* .:********** :**********:.*****::*
A1_3BTA_HC      TQEIKQRVVFKYSQMINISDYINRWIFVTITNNRLNNSKIYINGRLIDQK
E1_HC           NRGINQKLAFNYGNANGISDYINKWIFVTITNDRLGDSKLYINGNLIDQK
                .: *:*::.*:*.:  .******:********:**.:**:****.*****
A1_3BTA_HC      PISNLGNIHASNNIMFKLDGCRDTHRYIWIKYFNLFDKELNEKEIKDLYD
E1_HC           SILNLGNIHVSDNILFKIVNCSYT-RYIGIRYFNIFDKELDETEIQTLYS
                .* ******.*:**:**: .*  * *** *:***:*****:*.**: **.
A1_3BTA_HC      NQSNSGILKDFWGDYLQYDKPYYMLNLYDPNKYVDVNNVGIRGYMYLKGP
E1_HC           NEPNTNILKDFWGNYLLYDKEYYLLNVLKPNNFIDRRKDSTLSIN-----
                *:.*:.*******:** *** **:**: .**:::* .: .  .
A1_3BTA_HC      RGSVMTTNIYLNSSLYRGTKFIIKKYA-SGNKDNIVRNNDRVYINVVV-K
E1_HC           ---NIRSTILLANRLYSGIKVKIQRVNNSSTNDNLVRKNDQVYINFVASK
                    : :.* * . ** * *. *::   *..:**:**:**:****.*. *
A1_3BTA_HC      NKEYRLATNASQAGVEKILSALEIPDVGN-LSQVVVMKSKNDQGITNKCK
E1_HC           THLFPLYADTATTNKEK---TIKISSSGNRFNQVVVMNSV------GNCT
                .: : * :::: :. **   :::*.. ** :.*****:*       .:*.
A1_3BTA_HC      MNLQDNNGNDIGFIGFHQFNNIAKLVASNWYNRQIERSSRTLGCSWEFIP
E1_HC           MNFKNNNGNNIGLLGFKAD----TVVASTWYYTHMRDHTNSNGCFWNFIS
                **:::****:**::**:      .:***.**  ::.  :.: ** *:**.
A1_3BTA_HC      VDDGWGERPL
E1_HC           EEHGWQE--K
                 :.** *
```

**Table S1.** List of protonated residues in BoNT/A1 according to the studied system. The residues names correspond to: protonated histidine on N$\delta$ (HSD), protonated histidine on N$\epsilon$ (HSE), doubly protonated histidine (HSP), glutamate protonated on sidechain carboxyl (GLU), aspartate protonated on sidechain carboxyl (ASP). The Histidine located in the catalytic site of LC are written in bold. For each protonated residue the value of the mid-point pK(1/2) of the titration curve predicted by H++ is reported in parenthesis. In the majority of cases, the curve can be approximated by the classical sigmoidal (Henderson-Hasselbalch) shape, in which case pK(1/2) = pKa [3]. The protonation state of each titrable residue is then assigned by H++ basing on the comparison between pK(1/2) and the selected pH value. If pK(1/2) is $\geq$ pH the residue is considered as protonated.

| System | Residues |
|---|---|
| A1clo47 | LC: HSE-39(<0.0) ASP-58(6.6) GLU-64(5.1) GLU-147(8.6) GLU-148(8.5) GLU-164(5.6) HSP-170(6.3) GLU-197(5.8) GLU-279(6.9) **HSP-223**(>12.0) **HSP-227**(>12.0) **HSE-230**(<0.0) HSP-269(>12)<br>HC: ASP-442(4.8)<br>belt: ASP-513(5.8) GLU-508(4.8) GLU-519(9.1)<br>$HC_{NT}$: HSE-542(0.9) HSP-551(6.3) ASP-606(4.8) GLU-607(7.5) GLU-610(6.5) ASP-686(6.4) GLU-731(6.2) ASP-757(6.4) GLU-799(5.9)<br>$HC_{CN}$: GLU-874(5.8) HSP-877(6.8) HSP-1038(7.3) HSP-1054(6.0)<br>$HC_{CC}$: HSP-1243(5.3) GLU-1273(5.9) |
| A1clo47r | LC: HSD-39(<0.0) GLU-55(4.9) ASP-58(5.0) GLU-126(4.9) GLU-147(7.2) GLU-148 (9.6) HSE-170(3.9) ASP-203(6.2) **HSP-223**(>12) **HSP-227**(>12) **HSP-230**(6.1) GLU-252(8.9) HSP-269(8.4)<br>HC: GLU-470(5.1) GLU-482(6.0)<br>belt: ASP-513(4.8) GLU-519(9.1)<br>$HC_{NT}$: HSE-542(0.1) HSP-551(7.1) GLU-598(4.9) ASP-603(6.0) GLU-607(7.6) GLU-610(7.0) ASP-640(4.8) GLU-656(5.2) ASP-686(5.8) GLU-694(5.0) ASP-697(5.4) GLU-727(7.7) GLU-799(5.9) ASP-838(5.6)<br>$HC_{CN}$: HSP-877(5.9) GLU-929(4.7) HSP-1038(7.3) HSP-1054(6.3)<br>$HC_{CC}$: HSP-1243(5.7) GLU-1273(6.6) |
| A1clo70 | LC: HSE-39(<0.0) GLU-147(8.7) GLU-148(8.5) HSE-170(6.3) **HSP-223**(>12.0) **HSP-227**(>12.0) **HSE-230**(<0.0) HSP-269(>12.0)<br>belt: GLU-519(9.1)<br>$HC_{NT}$: HSE-542(0.9) HSE-551(6.3) GLU-607(7.5)<br>$HC_{CN}$: HSE-877(6.8) HSP-1038(7.3) HSE-1054(5.9)<br>$HC_{CC}$: HSE-1243(5.3) |
| A1clo70r | LC: HSD-39(<0.0) GLU-147(7.2) GLU-148(9.6) HSE-170(3.9) **HSP-223**(>12.0) **HSP-227**(>12.0) **HSE-230**(6.1) GLU-252(8.9) HSP-269(8.4)<br>belt: GLU-519(9.1)<br>$HC_{NT}$: HSE-542(0.1) HSP-551(7.1) GLU-607(7.6) GLU-727(7.7)<br>$HC_{CN}$: HSE-877(5.9) HSP-1038(7.3) HSE-1054(6.3)<br>$HC_{CC}$: HSE-1243(5.7) |
| A1ope47 | LC: HSP-39(6.2) GLU-47(4.9) GLU-96(4.8) GLU-126(5.1) HSE-170(2.9) **HSP-223**(10.9) **HSP-227**(>12.0) **HSP-230**(5.6) GLU-252(7.0) HSE-269(<0.0)<br>$HC_{NT}$: GLU-478(6.2) GLU-481(8.7) GLU-503(5.0) GLU-508(5.8) GLU-525(5.2) HSE-542(1.0) HSP-551(6.4) GLU-607(7.6) GLU-746(4.9) GLU-747(5.0) GLU-748(5.4) GLU-799(9.5)<br>$HC_{CN}$: HSP-877(10.9) ASP-838(5.3) HSP-1038(6.4) HSP-1054(6.1)<br>$HC_{CC}$: ASP-1236(5.5) HSP-1243(5.9) |
| A1ope70 | LC: HSE-39(6.4) HSE-170(3.2) **HSP-223**(11.4) **HSP-227**(>12.0) **HSE-230** GLU-252(7.8) HSE-269(<0.0)<br>HC: GLU-481(8.7)<br>$HC_{NT}$: HSE-542(<0.0) HSE-551(6.4) GLU-607(7.4) GLU-799(9.6)<br>$HC_{CN}$: HSE-877(6.5) HSE-1038(6.1) HSE-1054(6.1)<br>$HC_{CC}$: HSE-1243(6.0) |

 

**Table S2.** List of protonated residues in BoNT/E1 according to the studied system. The definitions of residue names are the same than in Table S1. The Histidine located in the catalytic site of LC are written in bold. For each protonated residue the value of the mid-point pK(1/2) of the titration curve predicted by H++ is reported in parenthesis.

| System | Residues |
|---|---|
| E1clo47 | LC: HSP-56(6.7) HSP-125(6.6) ASP-142(4.9) GLU-159(6.6) HSP-176(6.0) **HSP-212**(>12.0) **HSP-216**(7.0) **HSP-219**(>12.0)<br>HC: GLU-450(5.6)<br>belt: GLU-475(5.0) HSP-509(8.0)<br>$HC_{NT}$: ASP-521(4.8) GLU-529(6.0) ASP-579(4.8) GLU-583(6.6) GLU-636(4.7) GLU-677(8.5) GLU-730(5.0) GLU-741(5.4) HSP-797(6.3) ASP-829(5.0)<br>$HC_{CN}$: HSP-953(5.2) GLU-961(6.2) HSP-1021(8.9)<br>$HC_{CC}$: HSP-1155(5.9) HSP-1224(5.8) HSP-1228(6.8) HSP-1243(5.3) |
| E1clo70 | LC: HSE-56(6.6) HSE-125(6.6) HSE-176(6.0) **HSP-212**(>12.0) **HSP-216**(7.0) **HSP-219**(>12.0)<br>belt: HSP-509(8.0)<br>$HC_{NT}$: GLU-677(8.5) HSE-797(6.3)<br>$HC_{CN}$: HSE-953(5.2) HSP-1021(8.9)<br>$HC_{CC}$: HSE-1155(5.9) HSE-1224(5.7) HSE-1228(6.8) HSE-1243(5.3) |
| E1ope47 | LC: HSP-56(6.8) HSP-125(7.4) GLU-133(7.0) HSE-176(0.6) **HSP-212**(9.6) **HSP-216**(>12.0) **HSP-219**(>12.0) GLU-304(6.5)<br>HC: GLU-430(5.0) ASP-443(5.2)<br>belt: ASP-482(4.9) HSP-509(7.5)<br>$HC_{NT}$: GLU-513(5.7) ASP-521(6.5) ASP-579(5.2) GLU-612(4.9) GLU-623(5.1) GLU-632(5.6) GLU-670(6.3) GLU-673(5.8) GLU-717(4.9) GLU-726(5.0) GLU-727(4.8) GLU-730(4.9) GLU-743(5.0) GLU-773(5.5) GLU-781(4.8) HSE-797(4.3) GLU-803(5.0) ASP-829(6.2)<br>$HC_{CN}$: ASP-866(7.2) GLU-894(4.7) HSE-953(3.0) GLU-955(5.6) GLU-961(4.7) HSP-1021(6.6) ASP-1024(6.7) GLU-1050(5.4)<br>$HC_{CC}$: HSE-1155(2.7) GLU-1169(5.4) HSP-1224(5.8) HSP-1228(6.8) HSP-1243(6.4) |
| E1ope47r | LC: HSP-56(6.6) GLU-106(7.3) HSP-125(8.8) GLU-133(9.9) GLU-154(5.9) HSE-176(<0.0) GLU-189(5.3) **HSP-212**(10.4) **HSP-216**(>12.0) **HSP-219**(>12.0)<br>HC: GLU-430(4.8) ASP-442(6.5)<br>belt: ASP-482(6.2) HSP-509(9.0) ASP-510(7.3)<br>$HC_{NT}$: GLU-513(6.3) GLU-544(5.3) GLU-583(6.6) GLU-670(6.3) GLU-726(5.1) GLU-730(5.9) GLU-773(5.3) GLU-781(4.9) HSP-797(5.7) GLU-803(4.9) ASP-829(4.9)<br>$HC_{CN}$: ASP-866(8.9) HSP-953(6.0) GLU-955(5.0) GLU-961(4.9) HSP-1021(7.5) ASP-1024(5.7) GLU-1050(6.2)<br>$HC_{CC}$: HSE-1155(4.2) GLU-1169(6.1) HSP-1224(4.8) HSP-1228(6.6) HSP-1243(6.3) |
| E1ope70 | LC: HSE-56(6.8) HSP-125(7.4) GLU-133(7.0) HSE-176(0.6) **HSP-212**(9.5) **HSP-216**(>12.0) **HSP-219**(>12.0)<br>belt: HSP-509(7.5)<br>$HC_{NT}$: HSE-797(4.3)<br>$HC_{CN}$: ASP-866(7.2) HSE-953(3.0) HSE-1021(6.6)<br>$HC_{CC}$: HSE-1155(2.7) HSE-1224(5.8) HSE-1228(6.8) HSE-1243(6.4) |
| E1ope70r | LC: HSE-56(6.6) GLU-106(7.2) HSP-125(8.8) GLU-133(9.9) HSE-176 **HSP-212**(10.5) **HSP-216**(>12.0) **HSP-219**(>12.0)<br>belt: HSP-509(9.0) ASP-510(7.3)<br>$HC_{NT}$: HSE-797(5.7)<br>$HC_{CN}$: ASP-866(9.0) HSE-953(6.0) HSP-1021(7.5)<br>$HC_{CC}$: HSE-1155(4.2) HSE-1224(4.9) HSE-1228(6.6) HSE-1243(6.3) |

**Table S3.** Restraints applied during the Modeller run in the closed state of A1 and in the open state of E1 to produce respectively the systems A1clo47r, A1clo70r and E1ope47r, E1ope70r. The atoms belonging to the belt $\alpha$ helix are written in bold.

| Restraints for A1 | | | |
|---|---|---|---|
| first atom | second atom | target distance (Å) | interval (Å) |
| **I$^{489}$-C$\gamma$1** | F$^{357}$-C$\gamma$ | 4.0 | 1.0 |
| **I$^{489}$-C$\gamma$1** | L$^{103}$-C$\delta$1 | 4.0 | 1.0 |
| **I$^{489}$-C$\gamma$1** | I$^{348}$-C$\gamma$1 | 4.0 | 1.0 |
| **Y$^{492}$-C$\gamma$** | F$^{498}$-C$\gamma$ | 4.0 | 1.0 |
| **Y$^{492}$-C$\gamma$** | I$^{348}$-C$\gamma$1 | 4.0 | 1.0 |
| **L$^{494}$-C$\delta$1** | I$^{1086}$-C$\gamma$1 | 4.0 | 1.0 |
| **Q$^{490}$-N$\epsilon$2** | S$^{875}$-N | 3.0 | 0.5 |
| K$^{343}$-N$\zeta$ | **D$^{487}$-O** | 3.0 | 0.5 |
| **Q$^{491}$-N$\epsilon$2** | N$^{1083}$-N$\delta$2 | 3.0 | 0.5 |
| S$^{110}$-N | **Y$^{492}$-OH** | 3.0 | 0.5 |
| S$^{110}$-O$\gamma$ | **Y$^{492}$-OH** | 3.0 | 0.5 |
| **Y$^{493}$-OH** | D$^{1161}$-O$\delta$1 | 3.0 | 0.5 |
| **Y$^{493}$-OH** | D$^{102}$-O$\delta$2 | 3.0 | 0.5 |
| **Q$^{491}$-N$\epsilon$2** | T$^{495}$-O$\gamma$1 | 3.0 | 0.5 |
| Restraints for E1 | | | |
| first atom | second atom | target distance (Å) | interval (Å) |
| **D$^{466}$-O** | L$^{470}$-N | 3.0 | 0.5 |
| **L$^{465}$-O** | I$^{469}$-N | 3.0 | 0.5 |
| **I$^{469}$-C$\gamma$1** | L$^{98}$-C$\delta$1 | 4.0 | 1.0 |
| **F$^{472}$-C$\gamma$** | I$^{102}$-C$\gamma$1 | 4.0 | 1.0 |
| **V$^{468}$-C$\gamma$1** | K$^{330}$-C$\delta$ | 4.0 | 1.0 |
| **L$^{465}$-C$\delta$1** | N$^{459}$-C$\gamma$ | 4.0 | 1.0 |
| **L$^{465}$-C$\delta$1** | **Y$^{461}$-C$\gamma$** | 4.0 | 1.0 |
| K$^{342}$-N$\zeta$ | **N$^{463}$-N$\delta$2** | 3.0 | 0.5 |
| K$^{342}$-N$\zeta$ | **N$^{466}$-O$\delta$1** | 3.0 | 0.5 |
| K$^{329}$-N$\zeta$ | **Q$^{467}$-N$\epsilon$2** | 3.0 | 0.5 |
| D$^{326}$-O$\delta$1 | **N$^{471}$-N$\delta$2** | 3.0 | 0.5 |
| D$^{326}$-O$\delta$2 | **N$^{471}$-N$\delta$2** | 3.0 | 0.5 |
| K$^{330}$-N$\zeta$ | **N$^{471}$-N$\delta$2** | 3.0 | 0.5 |

**Table S4.** Quality parameters for the starting points of simulations as well as for the PDB templates. The scores were calculated using Molprobity [4] and QMEAN [5] on the Web servers molprobity.biochem.duke.edu and swissmodel.expasy.org/qmean.

| BoNT type | conformational state | restraints Table S3 | qmean4 | qmean6 | Molprobity |
|---|---|---|---|---|---|
| A1 | closed trans model | no | -5.28 | -4.487 | 3.28 |
| A1 | closed trans model | yes | -5.47 | -4.77 | 3.32 |
| A1 | open cleaved X-ray | no | -3.43 | -2.98 | 2.77 |
| E1 | closed cleaved X-ray | no | -2.95 | -2.80 | 2.66 |
| E1 | open trans model | no | -4.82 | -4.01 | 3.32 |
| E1 | open trans model | yes | -5.06 | -4.32 | 3.18 |
| 3BTA (A1) | open | - | -5.70 | -4.61 | 3.47 |
| 3FFZ (E1) | closed | - | -4.46 | -3.93 | 3.27 |
